# Characteristics and Impact of the rNST GABA Network on Neural and Behavioral Taste Responses

**DOI:** 10.1101/2022.05.08.491089

**Authors:** Susan P. Travers, B. Kalyanasundar, Joseph Breza, Grace Houser, Charlotte Klimovich, Joseph Travers

## Abstract

The rostral nucleus of the solitary tract (rNST), the initial CNS site for processing gustatory information, is comprised of two major cell types, glutamatergic excitatory and GABAergic inhibitory neurons. Many investigators have described taste responses of rNST neurons, but the phenotypes of these cells were unknown. The current investigation used mice expressing ChR2 under the control of GAD65, a synthetic enzyme for GABA. *In vivo* single-unit recording of rNST cells during optogenetic stimulation allowed us to address two important questions: (1) what are the gustatory response characteristics of “optotagged”, putative GABAergic (G+_TASTE_) neurons? and (2) how does optogenetic activation of the rNST GABA network impact taste responses in non-GABAergic (G-_TASTE_) neurons? We observed that chemosensitive profiles of G+_TASTE_ neurons were similar to non-GABA taste neurons but had much lower response rates. We further observed that there was a population of GABA cells unresponsive to taste stimulation (G+_UNR_) and located more ventrally in the nucleus. Activating rNST inhibitory circuitry suppressed gustatory responses of G-_TASTE_ neurons across all qualities and types of chemosensitive neurons. Tuning curves were modestly sharpened but the overall shape of response profiles and the ensemble pattern remained highly stable. These neurophysiological effects were consistent with the behavioral consequences of activating GAD65-expressing inhibitory neurons using DREADDs. In a brief-access licking task, concentration-response curves to both palatable (sucrose, maltrin) and unpalatable (quinine) stimuli were shifted to the right when GABA neurons were activated. Thus, the rNST GABAergic network is poised to modulate taste intensity across the qualitative and hedonic spectrum.

**SIGNIFICANCE STATEMENT:** The rNST, the CNS gateway for taste, is rich in GABAergic neurons and synapses. Our *in vivo* recordings from GAD65/ChR2 mice reveal that gustatory response profiles of optotagged GABAergic neurons resemble non-GABAergic neurons, but with much reduced amplitudes. A novel population of GABA neurons were unresponsive to oral stimulation suggesting they are targets for centrifugal influences. Activating rNST inhibitory circuitry modestly sharpened gustatory tuning but preserved the ensemble pattern for taste quality despite markedly suppressed responses. In behaving mice, activating rNST GAD65-expressing neurons with DREADDs shifted response-concentration curves for palatable and unpalatable stimuli, but preserved appropriate behaviors. These observations unveil previously unknown features of rNST GABA cells and demonstrate substantial inhibitory modulation at the first central taste relay.

## Introduction

The rostral nucleus of the solitary tract (rNST) is the initial CNS site for processing gustatory information. This processing relies not only on taste afferent input transmitted by the VIIth and the IXth cranial nerves, but also synaptic interactions between constitutive neurons and influences from extrinsic sources. There are two major types of rNST neurons: excitatory, glutamatergic neurons that often project outside the NST, including the parabrachial nucleus (PBN) (Gill et al., 1999), and GABAergic inhibitory neurons, many comprising local interneurons (Lasiter and Kachele, 1988; Davis, 1993). These generalizations are tempered by recent reports that some GABA neurons may have long-range projections (Shi et al., 2021) and the possibility of glutamatergic interneurons (McDougall et al., 2009; Corson and Bradley, 2013).

The gustatory response properties of glutamatergic and GABA neurons have not been directly assessed. Numerous investigators have reported single-neuron rNST taste responses but to our knowledge, in all cases the neurotransmitter phenotype of the cells was unknown. However, some insights can be gleaned from studies comparing rNST cells projecting to PBN (presumed glutamatergic) versus those that do not (presumed GABAergic). Based on antidromic activation, both activated and non-activated cells respond to a range of taste qualities but a consistent distinction is that PBN projection cells are activated more vigorously (Ogawa et al., 1984; Monroe and Di Lorenzo, 1995; Cho et al., 2002; Geran and Travers, 2009). In these studies, however, the identity of the non-antidromically activated neurons was ambiguous. Most importantly, whether these cells were GABAergic is uncertain since they may have been a different interneuron type or have projected to other rNST targets such as the reticular formation or caudal, visceral NST (Travers and Travers, 2018). In the present report we used transgenic mice expressing Channelrhodopsin-2 (ChR2) under the control of GAD65, a synthetic enzyme for GABA. This allowed us to use “optotagging” (Lima et al., 2009; Deubner et al., 2019) in an *in vivo* preparation to compare response characteristics of non-GABAergic and GABAergic taste neurons and to identify a novel population of NST GABA neurons unresponsive to taste stimulation.

The expression of ChR2 in GABAergic neurons further allowed us to evaluate how activating the rNST GABA network affected responses of non-GABA neurons. The rNST GABA network is comprised of synapses from local GABAergic interneurons as well as GABA terminals from the cNST (Travers et al., 2018) and central nucleus of the amygdala (Saha et al., 2000; Saha et al., 2002; Bartonjo and Lundy, 2020; Jin et al., 2021). Using the same mouse model and an *in vitro* slice preparation, optogenetic activation of the local rNST network suppressed rNST responses to afferent stimulation (Chen et al., 2016), consistent with suppressive effects of infusing GABA agonists *in vitro* (Wang and Bradley, 1993) or *in vivo* (Smith and Li, 1998).

In the visual cortex, optogenetic approaches have demonstrated that GABA interneurons have diverse effects on sensory responses categorized as “divisive”, mainly impacting response gain, or “subtractive”, sharpening tuning (Wilson et al., 2012). *In vivo*, infusion of GABA_A_ receptor antagonists into the NST increased gustatory responsiveness and resulted in broader tuning profiles (Smith and Li, 1998), suggesting that tonic activity in rNST GABA neurons exerts subtractive effects. In contrast, a previous *in vitro* study in our lab suggested that the major impact of the optogenetic release of GABA on afferent responses was divisive not subtractive (Chen et al., 2016). In the current experiment, we further explored the impact of optogenetic activation of the GABA network in the gustatory NST *in vivo* and demonstrate a modest but significant sharpening of response profiles coincident with marked suppression. At the population level, these effects gave rise to a stable representation in the ensemble code for taste quality. The widespread suppressive effects across taste qualities were echoed in behavioral effects of activating local GAD65-expressing neurons with DREADDs, which shifted the response-concentration curves for both palatable and unpalatable stimuli to the right in a brief-access licking task.

## MATERIALS and METHODS

### NEUROPHYSIOLOGY

#### Mice

All procedures involving animals were approved by the Ohio State University IACUC. Most mice expressed ChR2 and EYFP under the control of the GAD65 promoter (N = 66). These “GAD65 ChR2/EYFP” mice were generated by crossing a GAD65 cre line (Jax 010802) with mice carrying a floxed ChR2/EYFP allele (JAX 012569). This cross yielded expression of the fluorescent protein throughout the brain, including the entire NST. Expression was copious in the neuropil, but also evident in soma when inspected with confocal microscopy (**Figure 1A and B**). Three mice were generated from a cross of the same ChR2 mice with a strain expressing cre under the control of VGAT (Jax 028862). Thus, all mice expressed ChR2 in GABAergic inhibitory neurons. The strains were pooled for analysis since there were no obvious differences in response properties or effects between them. Both males and females were used (27 females, 39 males, 3 unknown) and the results from both sexes were combined in the analyses presented below since inspection of the data yielded no notable differences in the maximum response rates, the degree of inhibition or the proportion of taste responsive neurons directly driven by brain light between the two sexes (all P’s >.1 based on t-tests and chi-square evaluation). Mice were adults and ranged in age from 42-302 days (mean= 137.4 ± 7.2 days).

**Figure 1.**
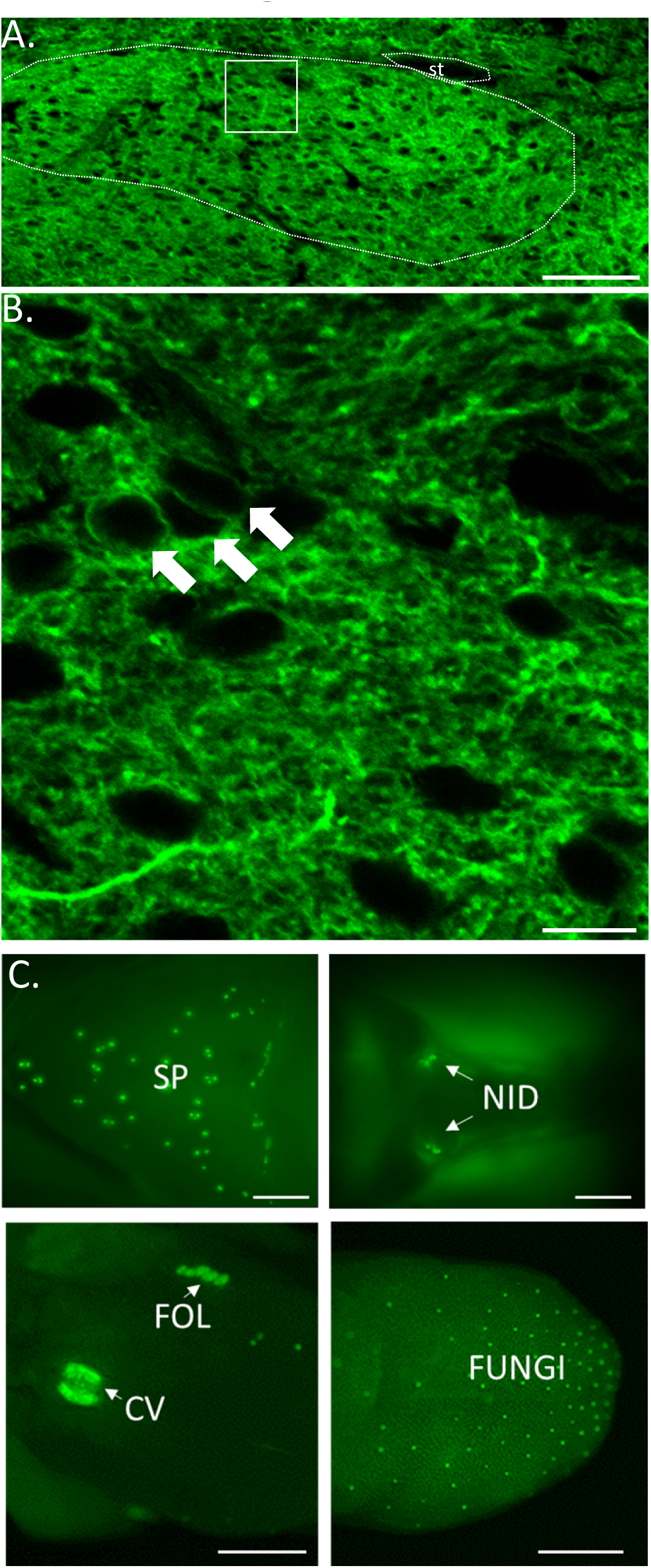
Expression of EYFP in the rNST and Taste Buds of a GAD65 ChR2/EYFP mouse. **A.** 20X confocal photomicrograph (single 2µm Z level) demonstrates EYFP expression throughout the nucleus. **B.** At a higher magnification (60X, digital zoom=3, single 1µm z level), EYFP expression is evident in somal membranes (arrows) and the neuropil. Scale bars: 100µm (A), 10µm (B). White dotted line indicates the approximate border of the nucleus. st-solitary tract. **C**. Low-power photomicrographs of native EYFP expression in different taste bud groups: soft palate (SP), nasoincisor ducts (NID), circumvallate (CV), foliate (FOL), and fungiform papillae (FUNGI). Scale bars = 1 mm.

### Surgical Preparation for Recording

To prepare mice for recording, animals were injected with urethane (1g/kg, I.P.). Throughout surgery, the level of anesthesia provided by this single dose of urethane was supplemented with isoflurane (<1% in O_2_) or sodium pentobarbital (∼25mg/kg) titrated to achieve an areflexive state. To view the mouth clearly and achieve optimal conditions for fluid and light stimulation, sutures were passed through the maxillary and mandibular lips for retraction, the hypoglossal nerves severed, and a tracheal cannula inserted (Breza and Travers, 2016; Kalyanasundar et al., 2020). We placed the animal in a stereotaxic device, made an incision over the skull and removed a portion of the interparietal plate with a drill and rongeurs to gain access to the rNST. A bolt was glued to the skull between bregma and lambda to stabilize the head without using the mouthpiece, further maximizing our view of the mouth. Following surgery and throughout recording, in most cases (N=63/66), mice were maintained in an areflexive state with isoflurane. In the remaining 3 subjects, anesthesia was supplemented by sodium pentobarbital or urethane.

### Taste Stimulation

Gustatory stimuli were made with chemicals purchased from Fisher or Sigma and diluted in artificial saliva (AS, in mM: 22 KCl; 15 NaCl, 0.3 CaCl_2_, 0.6 MgCl_2_) (Breza et al., 2010). Taste stimuli were delivered from pressurized glass bottles, connected via polyethylene tubes to a manifold (Warner Instruments) attached to a glass tube (1.0-1.2 mm) that was the final common path to deliver fluids to the mouth. In the initial one-third of preparations, the flow rate was ∼0.23 ml/s but we increased the rate to 0.6 ml/s in later preparations to make it easier to stimulate the whole mouth. Taste stimuli were applied for 10 s, preceded and followed by the flow of AS. The duration of the AS rinse was at least 20 s and a period of one minute or more separated successive taste stimulations. Taste stimuli representing the 5 classic taste qualities were used at mid-high to mid-range concentrations, referred to as stimulus set “A” and “B” respectively (**Table 1**); the five stimuli were presented in varied order. We began by using the lower concentrations but switched to the higher intensity stimuli because we were interested in evaluating whether inhibition sharpened tuning curves and the initial experiments with the mid-range stimuli revealed many narrowly-tuned units. Thus, we transitioned to the higher concentrations to attempt to broaden response profiles (Wu et al., 2015). In the results below, we use data from neurons tested with either stimulus set for describing general properties of the neurons and characteristics of putative inhibitory NST neurons. However, we restricted analyses to stimulus set A when describing effects of activating NST inhibitory circuitry on putative excitatory taste neurons, which involved more nuanced comparisons of the representation of different qualities under the two conditions. **Table 2** details the numbers of cells tested with each stimulus set.

**TABLE 1.** STIMULI

| QUALITY | CHEMICAL | ABBREVIATION | CONCENTRATIONS (mM) |  |
| --- | --- | --- | --- | --- |
|  |  |  | stimulus set A | stimulus set B |
| "sweet" | sucrose | SUC | 600 | 300 |
| "umami" | MSG +<br>2.5mM IMP +<br>100uM<br>amiloride | MSG<sub>ai</sub> | 600 | 600 |
| "salty" | NaCl | Na+ | 300 | 100 |
| "sour" | citric acid | CIT | 30 | 10 |
| "bitter" | cycloheximide<br>/<br>cycloheximide<br>+<br>quinine | BIT | 0.01 + 2.7 | 0.01 |

**TABLE 2:** Stimuli Used for Taste-Responsive Cells

| Neuron Type | Stimulus Set | # cells tested | Additional cells w/ 0.1M OR or no umami |
| --- | --- | --- | --- |
| **G<sup>-</sup>TASTE (N=84)** | A | **54\*** | 5 |
|  | B | 16 | 9 |
| **G<sup>+</sup>TASTE (N=12)** | A | 8 | 2 |
|  | B | 2 | 0 |

\* neurons used for analyses of effects of activating the inhibitory network on taste responses

### Neural Recording and Testing Protocol

#### Search Tracks

During search tracks for locating the gustatory NST, neural activity was recorded through epoxylite-coated tungsten microelectrodes ∼1-3 mΩ (Frederick Haer Inc. or World Precision Instruments). Signals were amplified and filtered, (10,000X; 600-10K; Alpha Omega, MCPplus), listened to using an audiomonitor (Grass AM8), and displayed on an oscilloscope and computer monitor using AD hardware and software (Cambridge Electronics Design [CED], Spike 2). We probed for taste-driven NST activity by flowing a mixture of taste stimuli (in mM: 300 sucrose, 10 citric acid, 100 NaCl, and 0.01 cycloheximide) and individual tastants over the entire oral cavity. Initial coordinates were typically 2.5mm caudal to lambda and 1.0mm lateral to the midline. Because the GAD65-ChR2 strain expresses ChR2 in taste buds throughout the oral cavity (**Figure 1C**), prominently in Type I (**Figure 1-1A**) (Baumer-Harrison et al., 2020), but perhaps also weakly in other taste bud cell types (Smith and Li, 1998; Larson et al., 2021), the search for the gustatory NST was facilitated by directing short pulses (5ms, ∼1-10Hz) of blue light (473 nm; 10 mW) to oral regions populated by taste buds using an LED source (Thor Labs, DC4104) controlled by CED hardware, CED software, a Grass S48 stimulator, and delivered though a 400 uM, 0.39 NA optical fiber (hereafter referred to as “light_m_”; i.e., light to the mouth). The light_m_ stimulus elicited robust responses in virtually all taste neurons; regardless of the quality they were responsive to. In a separate series of multiunit experiments using the GAD65-ChR2 mice we found that these responses were greatly attenuated by topical application of a P2X3 antagonist, AF343, suggesting that taste bud responses from the light_m_ stimulus activate primary afferent fibers requiring the same final common mechanism as natural tastants (**Figure 1-1B**). We also used blunt glass probes or brushes to stroke the oral tissues using non-nociceptive levels of force. Taste-responsive NST locations responded to light_m_ and sometimes to the somatosensory stimulus but NST sites purely responsive to the somatosensory stimulus were not activated by mouth light and were typically located lateral to the gustatory-responsive zone.

### Single-unit recording

#### Identifying Taste or Brain-light Responsive Cells

After locating the gustatory NST, we switched to recording using an “optrode”, a tungsten electrode (∼2-5 mΩ) combined with a 100 µm, 0.22 nA optical fiber (Thor Labs UM22-100). The optical fiber was configured to terminate < 1 mm dorsal to the electrode tip (mean distance = 660 ± 5.7 µm; range 550-820 µm, N = 110 measurements). To search for single units, we used taste stimuli and light_m_ and directed blue light pulses (5 ms, 1-10 Hz) to the NST through the optical fiber (hereafter also referred to as “light_br_”; i.e., light to the brain). Light pulses directed through the optrode were generated by a laser (Laserglow, LRS-0473-GFM-00050-05 or LRD-0470-PFFD-00100-05) and controlled by CED software and hardware. Because the laser does not deliver square pulses, measuring a constant light delivered though the optrode does not give an accurate measure of light intensity. Thus, we derived a better estimate using a power meter (Thor PM 100D) to measure the mean light intensity of each of a train of pulses and then averaged across the train. Across the 35 neurons for which this measure was made, pulses were 6.2 ± 0.2mW; the remaining neurons were stimulated using optrodes with similar configurations and laser settings. This brain light stimulus is expected to activate local GABAergic neurons, as well as fibers from GABA neurons outside the NST, notably the central nucleus of the amygdala (Saha et al., 2000; Saha et al., 2002; Bartonjo and Lundy, 2020; Jin et al., 2021) and the caudal NST (Travers et al., 2018). It is important to note that although GAD65 is expressed in just a subset of GABA neurons in some CNS locations such as the olfactory bulb (Parrish-Aungst et al., 2007), in the rNST it is highly co-localized with VGAT (Travers et al., 2018), suggesting that it is present in a majority of GABA (and glycinergic) neurons in this location.

### Testing

When a neuron was located that responded to taste (or mouth light), we screened it to determine if it also responded to brain light. Cells responding to light_br_ with short latency and in a time-locked, excitatory fashion presumably expressed ChR2 and were likely to be GABAergic. Therefore, we classified them as “G+_TASTE_” neurons; i.e., putative GABAergic taste-responsive neurons. Brain light stimulation consisted of a 10 Hz/10 s train of blue light pulses and/or a series of 20 pulses at 1, 4, 10, 20 and 50 Hz. G+_TASTE_ neurons were then tested with the standard protocol using the classic taste qualities (stimulus set A or B), presented in a varied order, to define their gustatory response profiles. Taste-responsive cells that did <u>not</u> respond in an excitatory, time-locked fashion to light were unlikely to be GABAergic and thus classified as non-GABA taste-responsive neurons (“G-_TASTE_”). This population is liable to include, though not be solely comprised of, glutamatergic neurons, including those that project to the parabrachial nucleus (Gill et al., 1999). G-_TASTE_ cells were tested with the panel of taste stimuli with and without concurrent stimulation with light_br_ (10 Hz) to determine effects of activating the NST GABAergic inhibitory network. Stimulations were repeated whenever possible. If the cell remained isolated, we used a micromanipulator to position the mouth light fiber so that it elicited optimal responses from the cell, and then tested light_m_ responses using nominal frequencies of 2, 5, 10, 20, and 50 Hz for 2 s (because frequency was manually controlled with the Grass Stimulator, actual frequencies deviated slightly: mean ± S.E.M. = 2.3 ± 0.1, 5.3 ± 0.1, 10.5 ± 0.2, 20.5 ± 0.2, 50.1 ± 0.1). As for taste responses, mouth lite responses were evaluated in the presence and absence of light_br_.

If a neuron was identified by being driven by light_br_, we first evaluated the characteristics of the response to brain light using the protocol above, then determined if it was taste responsive. If the cell responded to gustatory stimulation, the chemosensitive profile was determined. If a brain-light responsive neuron was not activated by taste stimulation and remained isolated, we tested it with oral somatosensory stimuli including gentle stroking of the oral tissues with a blunt glass probe and depressing the mandible. Brain-light responsive neurons unresponsive to taste were classified as “G+_UNR_” neurons (inhibitory unresponsive neurons), unless an oral somatosensory response could be identified, in which case they were deemed “G+_MECH_” (inhibitory mechanically-responsive) cells.

### Neurophysiological Data and Statistical Analysis

#### Basic Measures

##### Taste and Mouth Light Responses

Responses to both taste and light_m_ stimulation were calculated as evoked activity adjusted by a comparable period without stimulation. Net taste responses were the number of spikes during the 10 s stimulus delivery minus those during the 10 s pre-stimulus AS period, averaged across trials. The pre-stimulus period was calculated separately for control stimulations and those with concurrent brain light, since activating the NST inhibitory network often decreased the spontaneous rate of a cell. When stimuli were repeated, the mean of the trials was used. The mean and standard deviation (SD) of the spontaneous and pre-rinse AS activity were also calculated across trials for each neuron. The response criterion was a net evoked firing rate ≥ 1Hz (i.e., at least 10 spikes for 10 s) and 2.5 X the SD of the mean response to AS (Nishijo and Norgren, 1991; Geran and Travers, 2006; Kalyanasundar et al., 2020). However, two neurons that did not strictly meet this criterion were included. One was a bitter-responsive neuron with a clear but delayed response and another responded with a marginal increase in activity (0.9 Hz) with our standard whole-mouth stimulation protocol but responded robustly if stimuli were preferentially directed at the posterior mouth. Net responses to oral stimulation with light were calculated in a similar fashion by summing across the 2 s period of stimulation and adjusting for a comparable unstimulated period. However, in the case of light_m_ stimulation, the net response was adjusted by the spontaneous rate, since AS did not flow during the mouth light stimulation. The criteria for a response to mouth light stimulation was the same as that for a taste response, 2.5 X SD of the spontaneous rate.

##### Brain Light Responses

To analyze responses to brain light, we triggered off the beginning of the stimulus pulse and set a 10 ms search window for detecting a spike. Latency, jitter (SD of latency) and the proportion of trials evoking a spike were calculated for a given frequency and averaged across any repeated trials.

#### Additional Neurophysiological Analyses

##### G-_TASTE_ neurons

Statistical analyses were conducted and graphs prepared in Systat (v13), Excel (2016), and GraphPad Prism (v9.1.1). Comparisons between net taste responses under control conditions and during brain light stimulation (light_br_) were restricted to neurons tested with stimulus set A and compared using repeated measures or mixed ANOVAs, as appropriate. To identify similarities between response profiles, we used hierarchical cluster analysis based on responses to representatives of the five standard taste qualities, Pearson’s correlations, an average amalgamation schedule, and consulted the scree plot to determine the number of neuron groups. To evaluate across-neuron patterns of activity, Pearson’s correlations were calculated between stimuli across taste-responsive cells during the control period and the period with simultaneous brain light, and then plotted in a multidimensional scaling space. Breadth of tuning was evaluated using three measures: (1) the number of compounds eliciting a significant response in the cell, (2) the noise:signal ratio (N:S ratio = response to the 2nd-most effective stimulus/response to the most effective stimulus (Spector and Travers, 2005) and (3) entropy (Smith and Travers, 1979) calculated by the formula H = -1.43 ∑_i=1-5_ p_i_ log p_i_. where p_i_ is the proportion of the summed responses arising from a given stimulus. The first measure utilizes all responses but ignores response magnitude whereas the N:S ratio uses only two responses but takes relative magnitude into account. The entropy measure also takes response magnitude into account and utilizes all the responses. For calculating the entropy measure we substituted a very small value for zeros and responses that decreased below baseline, because the measure cannot accommodate zeros or negative numbers (Smith and Travers, 1979). When calculating the N:S ratio, we omitted responses to MSG_ai_ because, at the concentration employed (600mM), this stimulus strongly activates sugar/umami-responsive cells and amiloride-insensitive NaCl-sensitive neurons due to the multiple constituents of the compound (i.e., Na+ and glutamate); therefore calculating noise:signal for this stimulus is inappropriate. Finally, we evaluated effects of activating the GABA network by deriving a threshold linear function (Semyanov et al., 2004; Atallah et al., 2012; Wilson et al., 2012; Chen et al., 2016). For this analysis, responses of the neuron under control conditions were ordered from largest to smallest and the responses under both conditions normalized to the largest control response. The relationship between control and light-modulated responses was then subjected to linear regression. The resulting slope captures the proportional (divisive) effect of inhibition whereas the intercept provides a measure of the subtractive effects of inhibition. A line with a slope of 1 and a *y*-intercept of 0 indicates no effect of inhibition. A purely divisive effect of inhibition changes only the slope, while a purely subtractive effect yields a line with a slope of 1 but with a *y*-offset.

### Reconstruction of Recording Sites

#### Marking of recording sites

In selected instances, recording sites were marked with an electrolytic lesion made by passing current (3–8 µA for 3-10 s) at the site of recording in the NST. In other cases lesions were made in the reticular formation or vestibular nucleus dorsal or ventral to the cell. At the end of recording, the animal was injected with a lethal dose of anesthesia (80 mg/kg ketamine and 100 mg/kg xylazine or sodium pentobarbital (100mg/kg) and perfused through the left ventricle with phosphate-buffered saline (PBS), and 4% paraformaldehyde (in 0.1 M phosphate buffer) containing 1.4% L-lysine acetate and 0.2% sodium metaperiodate (McLean and Nakane, 1974). Afterwards, the brain was fixed overnight in 20% sucrose paraformaldehyde, blocked in the coronal plane and then stored in sucrose-phosphate buffer prior to sectioning.

#### Histology and immunohistochemistry

Coronal sections of the medulla (40-50 µm) were sectioned into two series and processed and mounted immediately or stored at –20°C in cryoprotectant until processing. One series was either left unstained to view in darkfield, stained with Weil (Berube et al., 1965) or (most frequently) black gold for delineating myelinated fibers (Schmued and Slikker, 1999). Double immunohistochemistry for P2X2 and NeuN was usually performed on the second series as described in Breza and Travers (Breza and Travers, 2016). Except when noted, processing was done at room temperature and the diluent and rinsing agent was PBS. Sections were rinsed before and after treatment with 1% sodium borohydride and 0.5% H_2_O_2_. Nonspecific binding was suppressed and tissue permeabilized using “blocking serum”, a mixture of 0.3% Triton, 3% bovine serum albumin and 7.5% donkey serum, before adding the primary antibodies (anti-P2X2, 1:10,000 and anti-NeuN, 1:1,000 [P2X2: rabbit polyclonal antibody against the intracellular COOH terminus of the P2X2 receptor, Alomone Labs, cat. no. APR003, RRID:AB_2040054; anti-NeuN: AB_2040054: mouse monoclonal antibody, Millipore MAB 377, RRID: AB_2298772). After primary antibody incubation (48–72 h, 4°C), sections were thoroughly rinsed and then treated with the secondary antibody (biotinylated anti-rabbit IgG, 1:500, Jackson ImmunoResearch Laboratories, Inc., RRID: AB_2337965) in blocking serum) for 1.5 h followed by an avidin-biotin mixture (Elite Kit, Vector, RRID: ABb_2336819) diluted in 0.1 M PB and 0.1% BSA. The chromagen reaction involved a 15 min pre-soak in 0.05% 3, 3-diaminobenzidine-HCl with 0.015% nickel ammonium sulfate, before adding H_2_O_2_ to achieve a concentration of 0.0015%. This reaction labeled P2X2 afferents dark brown to black. For labeling cell bodies with NeuN, we repeated the steps commencing with an anti-mouse secondary antibody incubation (biotinylated anti-mouse IgG, Jackson ImmunoResearch Laboratories, Inc., RRID: AB_2338570) but did the chromagen reaction sans nickel which resulted in light brown staining of soma. Tissue sections were examined under darkfield and brightfield optics.

We located the cell in the anterior-posterior dimension based on the lesion site relative to the caudal and rostral borders of the rNST and the morphology of the section. Because lesions were large relative to the dorsoventral dimension of the taste responsive area and because our observations suggested that the cell could be located anywhere from the center of the lesion to its ventral extent (most common), we relied on protocol notes taken during the experiment as the more accurate indication of dorsoventral location. This also allowed us to reconstruct cell location in the dorsoventral axis for more cells, since only a minority were marked with a lesion at the site of recording. During recording, the transition from the overlying vestibular nucleus was typically evident based on a notable decline in the amplitude of neural activity and the appearance of gustatory- and/or mouth-light driven responses. We noted this transition and the depth of responses every ∼25-50 µm thereafter. The dorsal-ventral location of a neuron based on these written records was only used as the electrode passed from dorsal to ventral, since positions of electrophysiological landmarks typically shifted dorsally as the electrode was retracted.

### BEHAVIORAL STUDIES WITH DREADDs

#### Mice

To study effects of activating rNST inhibitory neurons on taste-driven behavior, we used 8 mice that expressed the inhibitory DREADD receptor, hM3D(Gq) (Urban and Roth, 2015) and the fluorescent protein, mCherry, in rNST GAD65 neurons following viral injections (details below). We tested licking in response to different tastes in a brief-access paradigm. Five subjects were crosses between homozygous GAD65 cre males (Jax 010802; same line as used for the ChR2 neurophysiology crosses) and females expressing the Venus protein under the control of the VGAT promoter (RRID: ISMR_RBRC09645, (Wang et al., 2009)); 3 of these mice were also positive for the Venus protein. Two other subjects were the offspring of a cross between homozygous GAD65 cre mice. No difference in injection site size was apparent for mice heterozygous or homozygous for GAD65 cre. Five additional mice that were GAD65cre X VGAT/Venus crosses (1 Venus positive) received injections of a cre-dependent AAV virus that only expressed the mCherry protein. Qualitative inspection of the injection sites in the Venus-positive mice suggested that the vast majority of rNST cells expressing mCherry also expressed the Venus protein, confirming the specificity of the injections in inhibitory neurons.

#### Viral Injections

An anesthetic state was induced with a cocktail of xylazine and ketamine (2.5 and 25 mg/ml, 0.06 ml/20g bw, I.P.) and maintained with isoflurane (0.5-1%). The scalp was sanitized with alternating swipes of 70% ETOH and iodine, and a midline incision made. A small hole was drilled in the skull over the rNST and the dura removed. We made bilateral 50 nL pressure injections through glass pipettes (tip size = 40-50 µm) using a General Valve Picospritzer and monitored volume by observing the meniscus though the microscope equipped with a micrometer. rNST injection coordinates were determined by locating taste-responsive activity with a search electrode. We then replaced the electrode with the injection pipette filled with the virus and monitored neural activity through the injection pipette to assure accurate placement in the dorsal-ventral axis. The injection pipette was left in place for about 10 min after the injection was made. Mice in the inhibitory DREADD (“hM3D(Gq)/mC”) group were injected with pAAV2-hSyn-DIO-hM3D(Gq)-mCherry (ADDGENE #44361, titre: 1.8X10E^13^ GC/ml); mice in the mCherry viral control group (“mC”) received pAAV2-hSyn-DIO-mCherry (ADDGENE #50459, titre: 2.6X10E^13^ GC/ml). Mice recovered for 2 weeks before training. Mice were run in squads at different time points; (hM3D(Gq)/mC - 2 squads; mC - 1 squad).

#### Experimental Design: Behavioral Training and Testing

Training and testing took place in a standard brief-access testing apparatus (Davis MS160-Mouse; DiLog Instruments, Tallahassee, FL), “Davis Rig”), in which stimulus presentation and timing was computer-controlled. Mice licked different sapid solutions from bottles fitted with sipper tubes that were automatically moved into position. A shutter controlled the mouse’s access to the bottle. For the hM3D(Gq)/mC group, the experiment proceeded in four blocks (1) training, (2) quinine testing, (3 and 4) sucrose and maltrin testing (with sucrose and maltrin order counterbalanced across two squads). Stimuli were reagent grade and dissolved in distilled water: quinine (µM: 30, 100, 1000, 3000), sucrose (mM: 30, 100, 200, 300, 600, 1000), and the maltodextin, maltrin 580 (%: 1, 2, 8, 16, 24, 32; a generous gift of the Grain Processing Corporation, Muscatine, IA). Mice in the mC control group only underwent the training phase followed by testing with sucrose. For the hM3D(Gq)/mC subjects, quinine was tested prior to sucrose and maltrin because pilot studies suggested that mice were reluctant to sample the aversive stimulus after first experiencing the appetitive compounds. Training and quinine blocks were performed under ∼23 hour water deprivation and occurred on a daily basis with a break between the training and quinine testing. When evaluating appetitive stimuli, mice were tested under 23.5 hour food deprivation and testing was performed on alternate days.

Training began two weeks following the viral injection. During the first two training days, mice were placed in the rig for 30 minutes with the shutter open and a single bottle of distilled water available. On days 3 and 4, mice were permitted water access in discrete trials: a bottle moved into position, the shutter opened and remained open for 5 s after the first lick then closed for 10 s before opening again. There was no restriction on the number of trials mice could initiate in the 30 minute period. Day 5 was identical to days 3 and 4, with the addition of a CNO injection to check for any untoward effects. This same 5 s trial structure was used for the taste stimulus testing. Prior to each taste test block, the relevant stimulus was introduced in a single session without any injections. Subsequently, a given stimulus was tested 4 times, twice with saline injections (∼0.1ml/10g bw, I.P.) and twice with CNO (1.0mg/kg, in saline, I.P.). Injections were given 30 minutes before the test session. Injection order was counterbalanced across mice. For the hM3D(Gq/Mc) groups, an additional 30 minute water trial session with a CNO injection was given between blocks testing different stimuli to assess the possible development of effects on lick rate over time. Following the last test period, mice were perfused as in the neurophysiological experiments and brains cut into 40 µm sections. Sections were mounted onto slides and fluorescent photomicrographs taken of the injection site. All injections were found to be centered in rNST and mainly confined to the nucleus.

#### Behavioral and Statistical Analysis

Mice were omitted from analysis if they sampled each stimulus on fewer than 2 trials/session; this yielded final N’s of 7 mice each for quinine and sucrose, and 6 for maltrin for the DREADD-injected mice. There were 5 mice in the mC control group. Responses to quinine were quantified as the ratio of the number of licks to quinine relative to the average number of licks to water for a given test day. For sucrose and maltrin, responses were quantified as a standardized lick ratio (Glendinning et al., 2002) where we normalized the number of licks relative to the maximum possible licks based upon the modal ILI for a given session. These standardized measures controlled for the small changes in lick rate that occurred in the sucrose and maltrin testing phases but not in the earlier quinine testing phase. Analyses were performed as repeated measures ANOVAs with concentration and drug state as the independent variables. These were followed by Bonferroni-adjusted paired t-tests for each concentration. Statistics were performed in Systat (v13), Excel (2016), and GraphPad Prism (v9.1.1). The significance level was set at *p* < .05. Fits of the average curves were made using non-linear regression with the equation:

Y = Minimum Response + (Maximum Response-Minimum Response)/(1+10^((LogEC50-X)*Hill^ ^Slope)^.

## RESULTS

### Putative GABA NEURONS (N=38)

Thirty-eight cells were classified as putative GABA neurons because they responded faithfully and at short latency to light pulses delivered through the optrode. Fifteen responded to taste (G+_TASTE_) and four to oral somatosensory stimulation or depressing the mandible (G+_MECH_, two of these were slightly lateral or ventral to the NST borders). An additional 19 neurons failed to respond to either taste stimulation or fluid flow (G+_UNR_) and a lack of responsiveness to stroking the oral mucosa was further confirmed for 9 of these cells. Quantitative data on evoked responses to 10 Hz light stimulation were obtained for 31 light-responsive neurons and for 26 cells over an extended frequency range. The remaining 7 light-entrained neurons were cells with obvious synchronized responses to light pulses and histologically verified to be in the NST but with isolation insufficient for accurate quantification (N=4) or for which only protocol notes were taken (N=3); these cells were used only in the histological analyses.

**Figure 2** shows examples of neurons classified as putative taste G+_TASTE_ (A-C) or unresponsive G+_UNR_ (D-F) inhibitory cells based on their reliable short-latency, low-jitter responses to brain light and the presence or absence of a taste response. The G+_TASTE_ cell responded most robustly to the bitter mixture and well to NaCl (not shown) but, somewhat atypically, was silenced by sucrose stimulation. In addition, light evoked a clear increase above baseline and, reliably followed light pulses at 10 Hz (**2B**). Panel **2C** shows the mean latency, jitter, and for the light-driven responses across all the G+_TASTE_ neurons for which we tested a range of light_br_ stimulation frequencies (N=11). An example of a G+_UNR_ neuron is shown in panels **D-E**. This neuron lacked spontaneous activity and a taste response and also did not respond to fluid flow, stroking the oral mucosa, or depressing the mandible. Indeed, this cell was detected only because of its synchronous response to light_br_ stimulation as the electrode passed to the ventral region of the nucleus (**E**) where taste activity was waning. Panel **F** shows the average characteristics of the light-driven responses across all G+_UNR_ neurons for across light_br_ stimulation frequencies (N=11). Panel **G** presents individual data for frequency following across the 26 putative inhibitory cells for which data for both 1 and 10 Hz light_br_ stimulation were available. On average, these cells followed 69% ± 7% of light pulses at 10 Hz and 87 ± 5% at 1 Hz (*p =*.001, paired t-test). Thus, like “optotagged” cells originally described by Lima in auditory cortex, (Lima et al., 2009) faithful following was apparent at 1 Hz. There were 5 additional neurons for which only 10 Hz data was available, but that exhibited similar properties (following= 54.0 ± 0.2% at 10Hz). Thus, we classified all 31 cells as putative inhibitory cells. There were three other neurons (2 unresponsive to taste and 1 with an oral somatosensory response) that exhibited clear increases in firing in response to light_br_ stimulation but followed 1 or 10 Hz light pulses on only about 1/5 of the trials. We were unable to unambiguously classify these cells and so did not include them in our analyses, but it was interesting that two of them followed higher frequencies more faithfully than lower ones. We speculated that temporal summation may have compensated for low ChR2 expression in these cases or that these cells reflected complex network effects.

**Figure 2.**
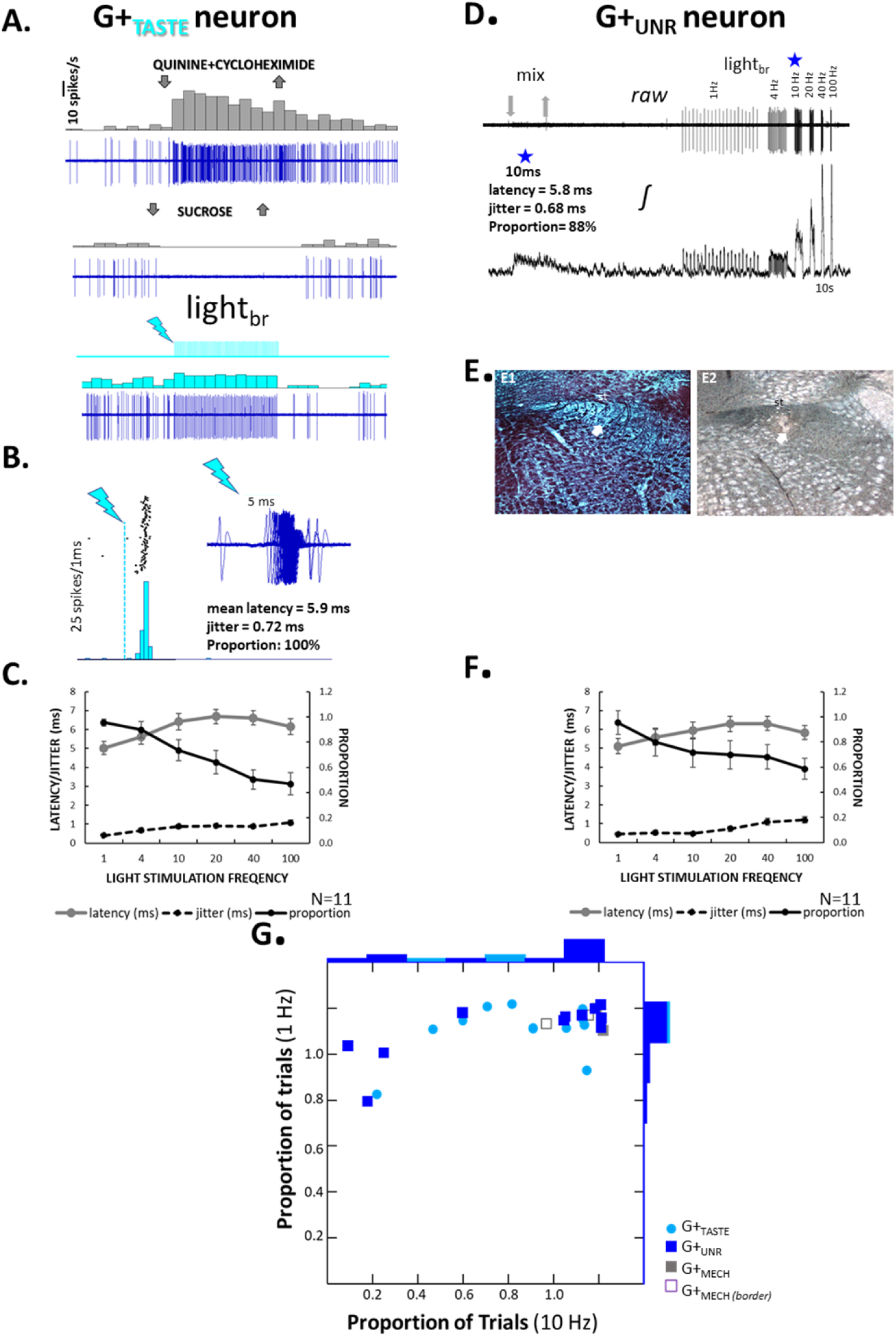
Some rNST putative inhibitory neurons are gustatory-responsive (G+_TASTE_, A-C) but others are unresponsive to orosensory stimulation (G+_UNR_, D-F). **A**. Response characteristics of an exemplar G+_TASTE_ neuron. Raw record and peri-stimulus time histograms (1 s bins) for a G+_TASTE_ neuron for the bitter mixture, sucrose and light_br_. **B**. Brain light-triggered histogram (1 ms bins) and triggered spikes (upper right); lightning bolts indicate the onset of the 5 ms light pulse. This neuron responded at a short latency with low jitter (note the sharp peak in the 5 ms bin in the histogram) and followed a 10 Hz pulse train faithfully (100% following at 10 Hz for 10 s). **C**. Average (± S.E.M) latency, jitter, and proportion of trials followed across the population of G+_TASTE_ cells. N= 15 for 10 Hz; N=10-11, other frequencies. **D-E**. Example of a G+_UNR_ neuron. **D**. Raw (top) and integrated (bottom) records showing recording from a non-spontaneously active neuron driven by light_br_ but unresponsive to taste stimulation. The raw record shows a miniscule background taste response which is more evident in the integrated record and was detectable on the audiomonitor. The cell followed light_br_ pulses reliably at 10 Hz. **E**. Post-hoc histology for the cell in Panel D (E1: brightfield image of black-gold staining; E2: darkfield image from an adjacent unstained section) indicates a location in the ventral region of the nucleus, corresponding to the ventral subdivision (Whitehead, 1986; Ganchrow et al., 2014). Scale bar = 100 µm. **F**. Average (± S.E.M) latency, jitter, and proportion of trials followed, across the population of G+_UNR_ cells for which we tested an extended frequency range (N=11). **G**. Frequency following characteristics for light_br_ stimulation at 1 and 10 Hz for all cells classified as putative GABA neurons (G+_TASTE_, G+_UNR_, and G+_MECH_) for which both frequencies were tested. At 1 Hz, each cell responded within 10 ms (mean latency = 4.93 ± .05 ms) on 73% or more of the trials.

Putative GABA and non-GABA neurons differed on the basis of characteristics other than their light-synchronized responses (**Figure 3**). Although many neurons in all three main cell groups had low spontaneous rates, the mean spontaneous rate was highest in the G-_TASTE_ cells and lowest in G+_UNR_ neurons (**Figure 3A**). Indeed, whereas the vast majority of G-_TASTE_ (94%) and all G+_TASTE_ neurons exhibited some level of resting activity, only 46% of G+_UNR_ neurons were observed to fire any spikes in the absence of light_br_ stimulation. In addition, based on microdrive coordinates, G-_TASTE,_ G+_TASTE,_ and G+_UNR_ neurons were preferentially staggered from dorsal to ventral (**Figure 3B**). This finding was consistent with what we observed after reconstructing the histology for the subset of cells for which we made lesions at the site of recording (**Figure 3C**.).

**Figure 3.**
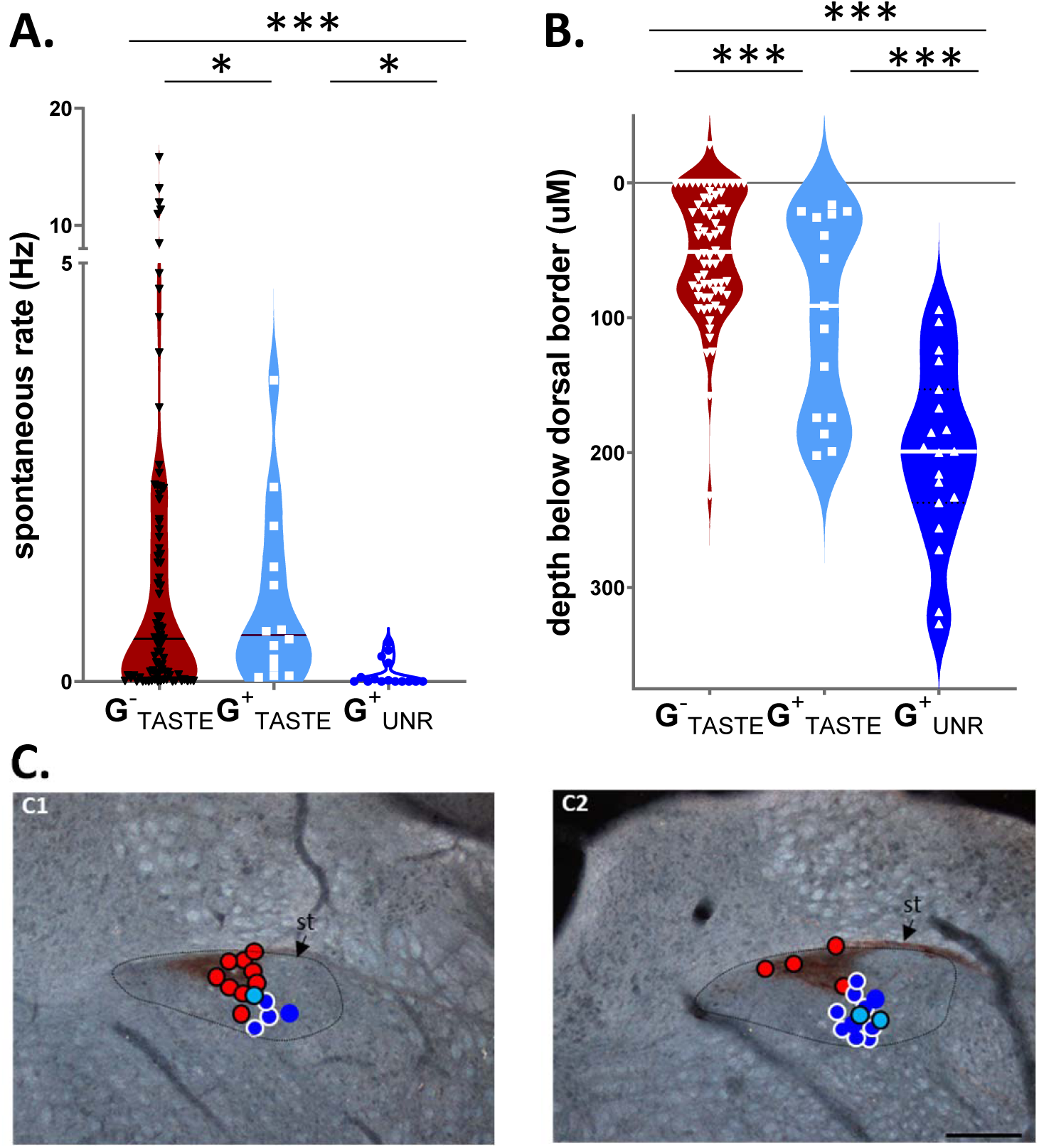
G-_TASTE_, G+_TASTE_ & G+_UNR_ neurons have distinctive distributions of spontaneous rate and locations in the dorsoventral axis. **A.** Violin plots depicting spontaneous rate of individual G-_TASTE_, G+_TASTE_ & G+_UNR_ neurons. Horizontal lines indicate medians. A Kruskal-Wallace ANOVA (N=113) indicated significant differences between all 3 cell types (***G^-^_TASTE_ vs G^+^_TASTE_, *p =*1.66e-06; G^-^_TASTE_ vs G^+^_UNR_, *p =* 1.56e-06, G^+^_TASTE_ vs G^+^_UNR_, *p =*1.13e-05). B. Depth of individual G-_TASTE_, G+_TASTE_ & G+_UNR_ neurons relative to the dorsal border of the NST identified electrophysiologically by the transition from high-amplitude activity characteristic of the vestibular nuclei to lower-amplitude activity responsive to gustatory and oral somatosensory stimulation. Horizontal lines indicate medians. ANOVA indicated a significant difference by type (*p =*1.75e-11, ANOVA, N=102) and Tukey’s post-hoc tests indicated significant differences *** between all 3 cell types (G^-^_TASTE_ vs G^+^_TASTE_, *p =*.008; G^-^_TASTE_ vs G^+^_UNR_, *p =*1.07e-05, G^+^_TASTE_vs G^+^_UNR_, *p =*1.17e-05). C. Locations of the subset of neurons marked by a lesion made at the site of recording; symbols are superimposed on darkfield photomicrographs of NST sections from more rostral (C1) and caudal (C2) levels of the rNST immunostained for P2X2 gustatory afferent fibers. Symbol colors are the same as for panel A, but outlines were added to enhance visibility: G-_TASTE_ neurons: red circles/black borders; G^+^_TASTE_ neurons: light blue circles/black borders, and G+_UNR_ neurons: dark blue circle/white borders. A few G+_MECH_ neurons are also plotted: blue circles/no outlines. st: solitary tract. Dotted outline indicates the approximate borders of the nucleus. Scale bar: 200 µm.

**Figure 4** summarizes the gustatory response characteristics of the population of putative inhibitory G+_TASTE_ cells that we were able to test with at least 4 of the 5 standard taste qualities (N=12) compared to “G-_TASTE_” (N=84). Stimuli representing each quality drove responses in both G+_TASTE_ and G-_TASTE_ cells (**4A**). Because there was only a small sample of G+_TASTE_ neurons, we did not consider it meaningful to compare response profiles of G+_TASTE_ and G-_TASTE_ neurons in detail. Thus, we did not use cluster analysis to divide them into chemosensitive groups or evaluate their breadth of tuning. However, as reflected in the overall mean responses, neurons optimally responsive to each taste quality were evident (**4B**), as was the case for G-_TASTE_ cells (see below). Despite these broad similarities, a salient characteristic of G+_TASTE_ neurons was that each taste stimulus (with the exception of the bitter mixture) elicited a markedly and significantly smaller response than in G-_TASTE_ cells. Indeed, even when the maximal response (MAX) for a given neuron was considered, the average taste-driven response of G+_TASTE_ cells was only 37% as great as in G-_TASTE_ neurons.

**Figure 4.**
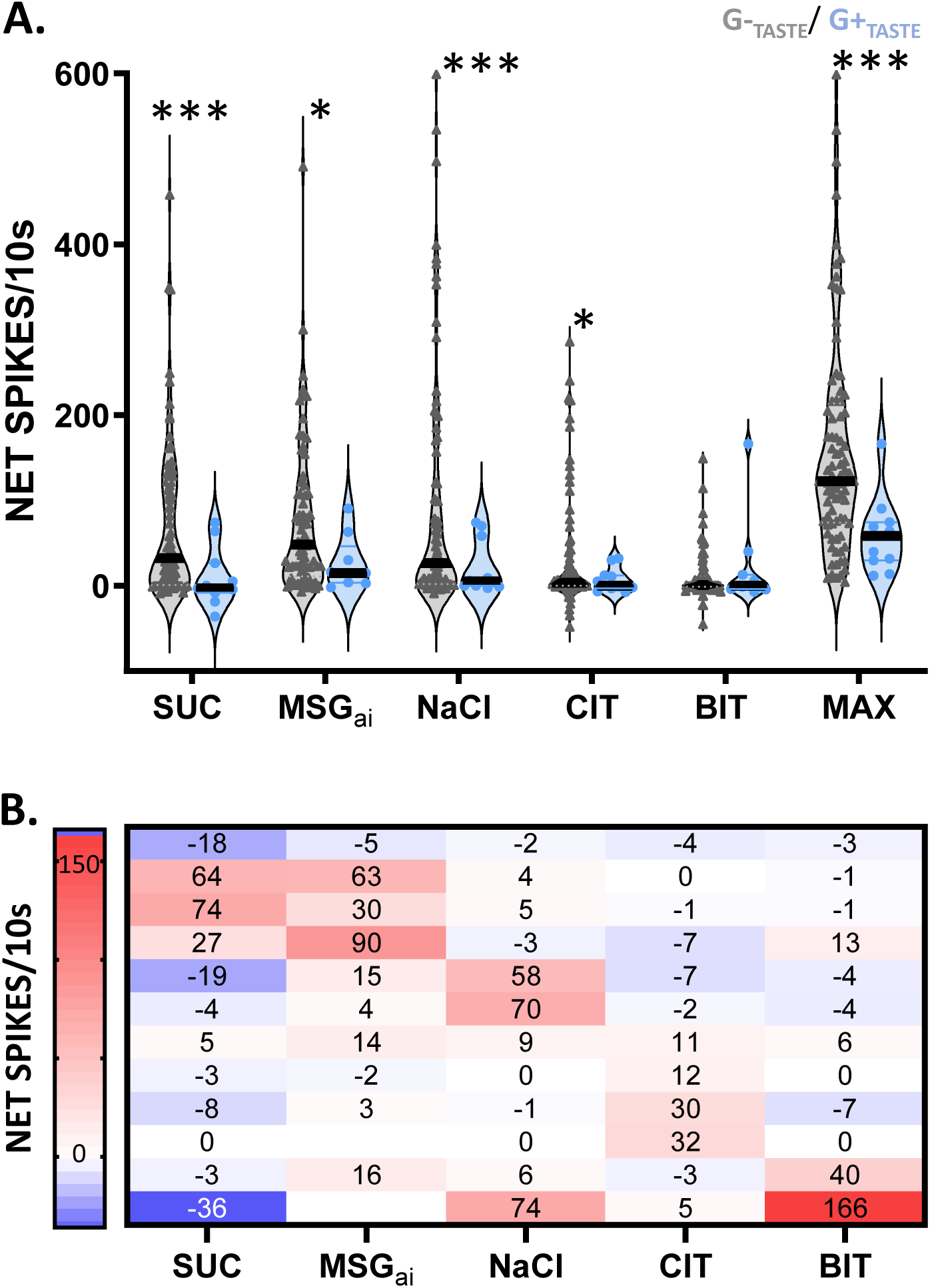
G-_TASTE_ and G+_TASTE_ neurons respond to the same taste qualities but responses are weaker in G^+^_TASTE_ cells. **A.** Violin plots showing the individual (symbols) and median (lines) responses to 5 standard taste stimuli and for the maximum response (MAX) for a given cell for G-_TASTE_ (N=84 except for BIT, N=82 and MSG_ai_, N=72) and G+_TASTE_ (N=11, except for MSG_ai_, N=9; one G+_TASTE_ neuron that only responded in an inhibitory fashion was omitted from these means but appears in the individual responses in panel B). An ANOVA (excluding MSG_ai_) showed a main effect for cell type: *p =*.009, but not stimulus: *p =*.118, or an interaction between stimulus and cell type (*p =*.147). Bonferroni-adjusted t-tests indicated that responses were significantly smaller for all stimuli except BIT (***SUC: *p =* 5.52e-04, MSGai: \**p =*.01, NaCl: \*\*\**p =*6.96e-04, CIT: \**p =*.01). The maximum response was also smaller (*p =*5.42e-6). **B.** Heat map with net responses showing individual response profiles for each G+_TASTE_ neuron.

Although G-_TASTE_ and G+_TASTE_ neurons were differentially distributed in the nucleus, there was clear intermingling between the various cell types. **Figure 5** shows an example where a light-driven G+_UNR_ neuron (larger potentials; blue wavemark) was recorded simultaneously with a G-_TASTE_ cell (small pink wavemark; only used for illustrative purposes). The G-_TASTE_ cell responded robustly to a mixture of the 4 classic taste stimuli (**5A**: mix) and to sucrose and MSG_ai_ but NaCl but citric acid and the bitter mixture were ineffective. Moreover, whereas activity in the G+_UNR_ neuron was synchronized to light_br_, the taste-elicited activity in the G-_TASTE_ neuron was suppressed (**5A**: mix + light_br_). The remainder of the paper considers such suppressive effects of activating the rNST GABA network on the taste responses of putative non-GABA cells.

**Figure 5.**
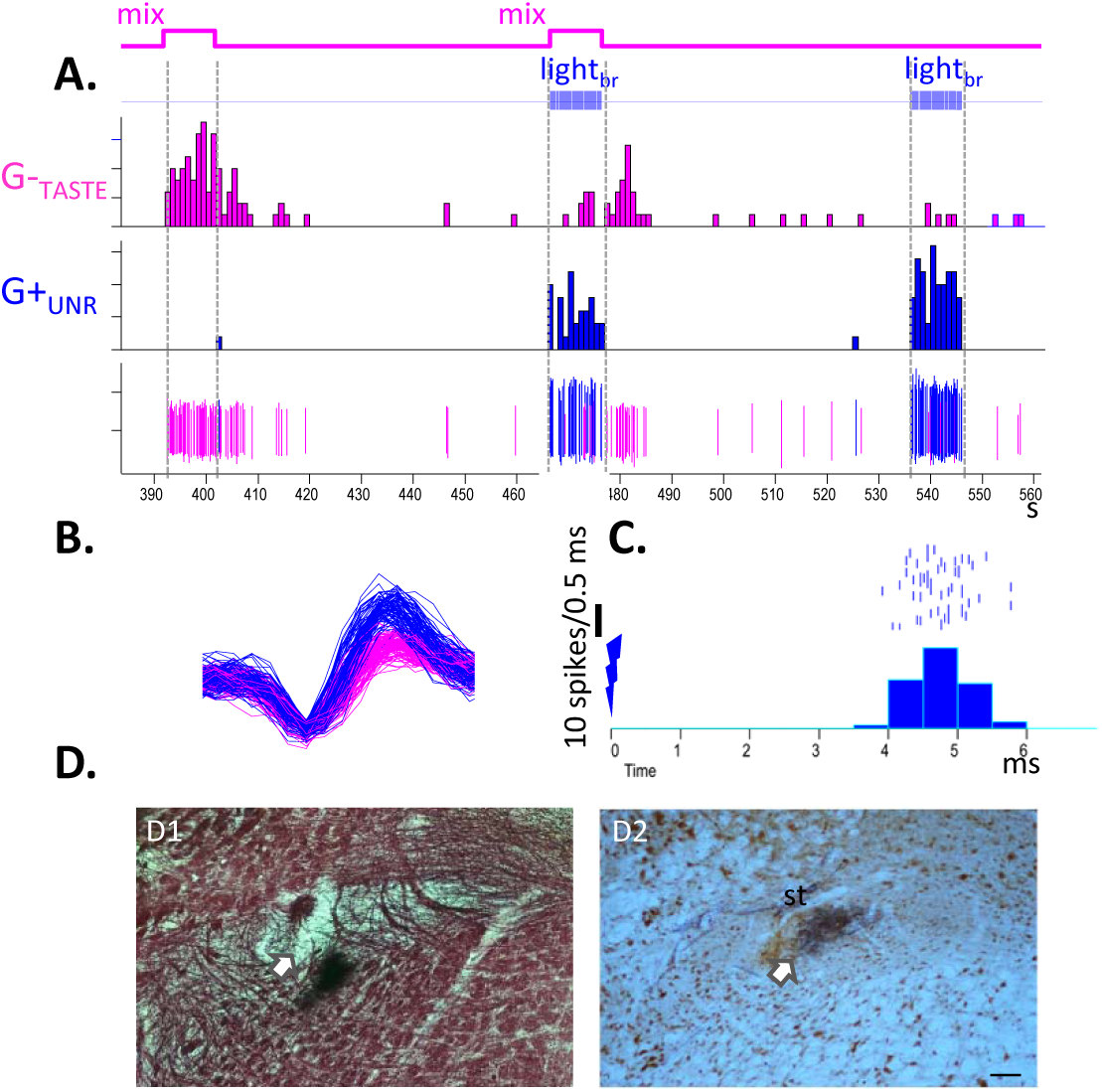
Some putative inhibitory rNST neurons are unresponsive to taste stimulation and intermixed with non-GABA taste responsive neurons. **A**. Peristimulus time histogram and windowed spikes showing a neuron unresponsive to gustatory stimulation but driven by light_br_ (i.e., a G^+^_UNR_ neuron, windowed in blue) recorded simultaneously with a (marginally isolated) taste-responsive neuron where light_br_ suppressed taste-elicited activity (i.e., a G^+^_TASTE_ neuron, windowed in pink). **B**. Expanded time base showing the waveform of the windowed cells. C. Light_br_ - triggered histogram (0.5 ms bins; light onset at lightning bolt) showing that the G^+^_UNR_ cell responded at a short latency (5.89 ms) and jitter (0.42 ms) on most (76/100 @ 10Hz) trials. D. Photomicrographs illustrating the lesion at the recording site (arrows). D1. Staining with black-gold showing myelinated fibers; D2 alternate sections were double immunostained for NeuN (brown) and P2X2 (black). Lesions were often evident based on homogeneous instead of cellular NeuN staining, presumably an immune reaction against damaged tissue due to the anti-mouse secondary antibody used to detect NeuN. Scale bar = 100µm.

### EFFECTS OF THE NST GABA NETWORK ON TASTE RESPONSES IN NON-GABA (G-_TASTE_) CELLS (N=54)

#### Population Effects

Across the 84 non-GABA taste cells described in the preceding section, 54 were tested with stimulus set A (Table 1) and used to analyze the effect of activating the rNST GABA network on a population which likely includes excitatory projection neurons. **Figure 6** shows that there was notable cell to cell variation in the degree of inhibition, whether assessed as absolute or percent suppression. The magnitude of the control response appears to be one factor underlying the variable suppression. Though it explains it only partially, the degree of absolute suppression was positively correlated with the size of the control response (Pearson’s r = 0.451, *p* =.001), whereas percent suppression was negatively correlated (r = -.432 *p* = .001). Thus, very small responses could be virtually obliterated during activation of the GABA network by eliminating a small number of spikes whereas, in highly-responsive neurons, more spikes were available to be suppressed but they represent a smaller fraction of the control response. Despite this cell-to-cell variability, optogenetic activation of the rNST GABA network substantially and significantly suppressed mean responses to each stimulus (**Figure 7A & B**), suggesting that entire gustatory spectrum is susceptible to inhibitory influences. The firing rate during sustained flow of artificial saliva likewise declined during light stimulation (11.8 ± 3.1 versus 7.5 ± 2.2 spikes/10 s, *p =*.001, N=54).

**Figure 6.**
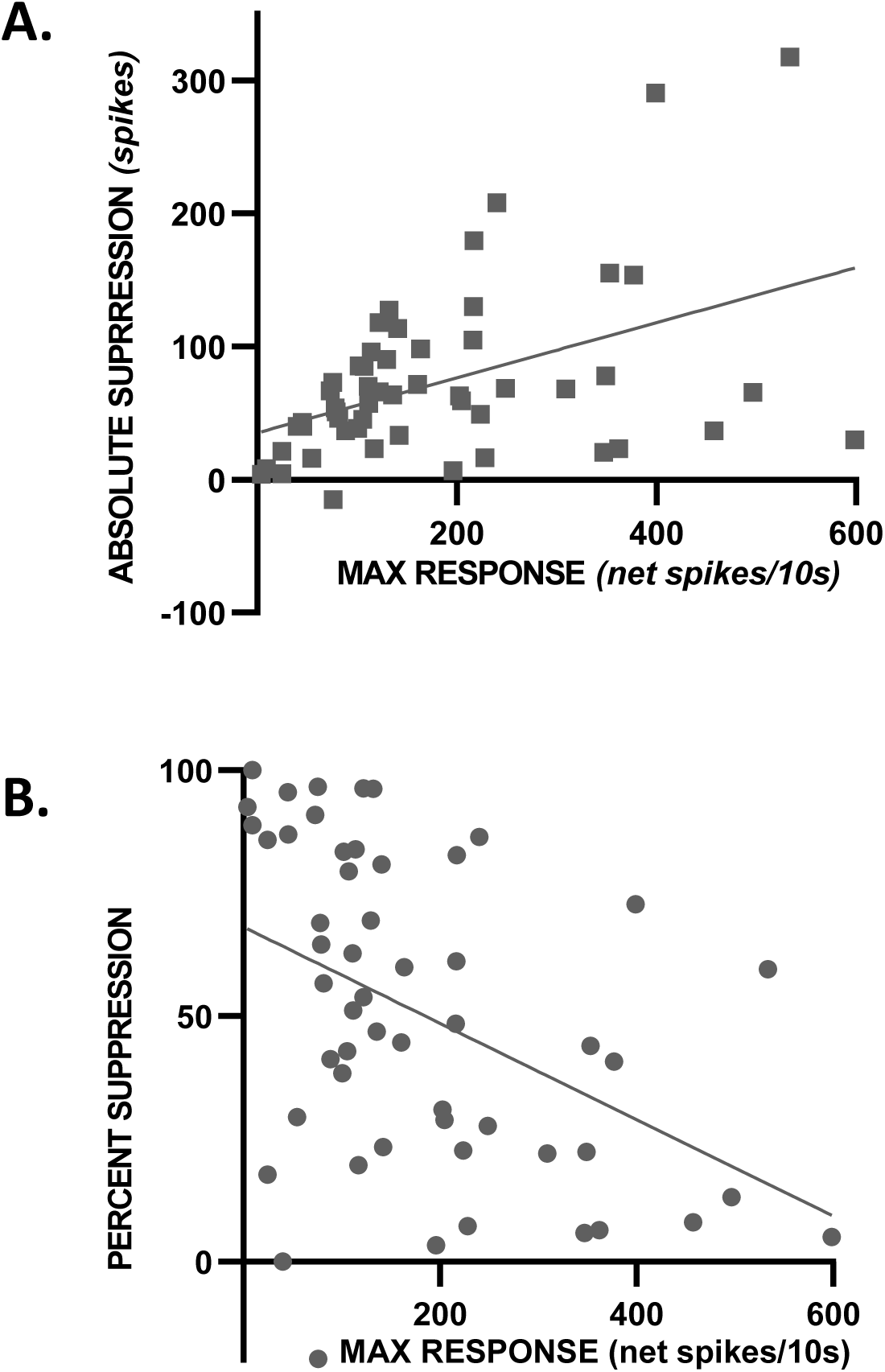
There is substantial cell-to-cell variation in the degree of GABA-network induced suppression. Scatterplot showing the degree of suppression induced by light_br_ stimulation expressed as absolute suppression (control response - light_br_ modulated response) and percent suppression ([control response-light_br_ modulated response]/control response).

**Figure 7.**
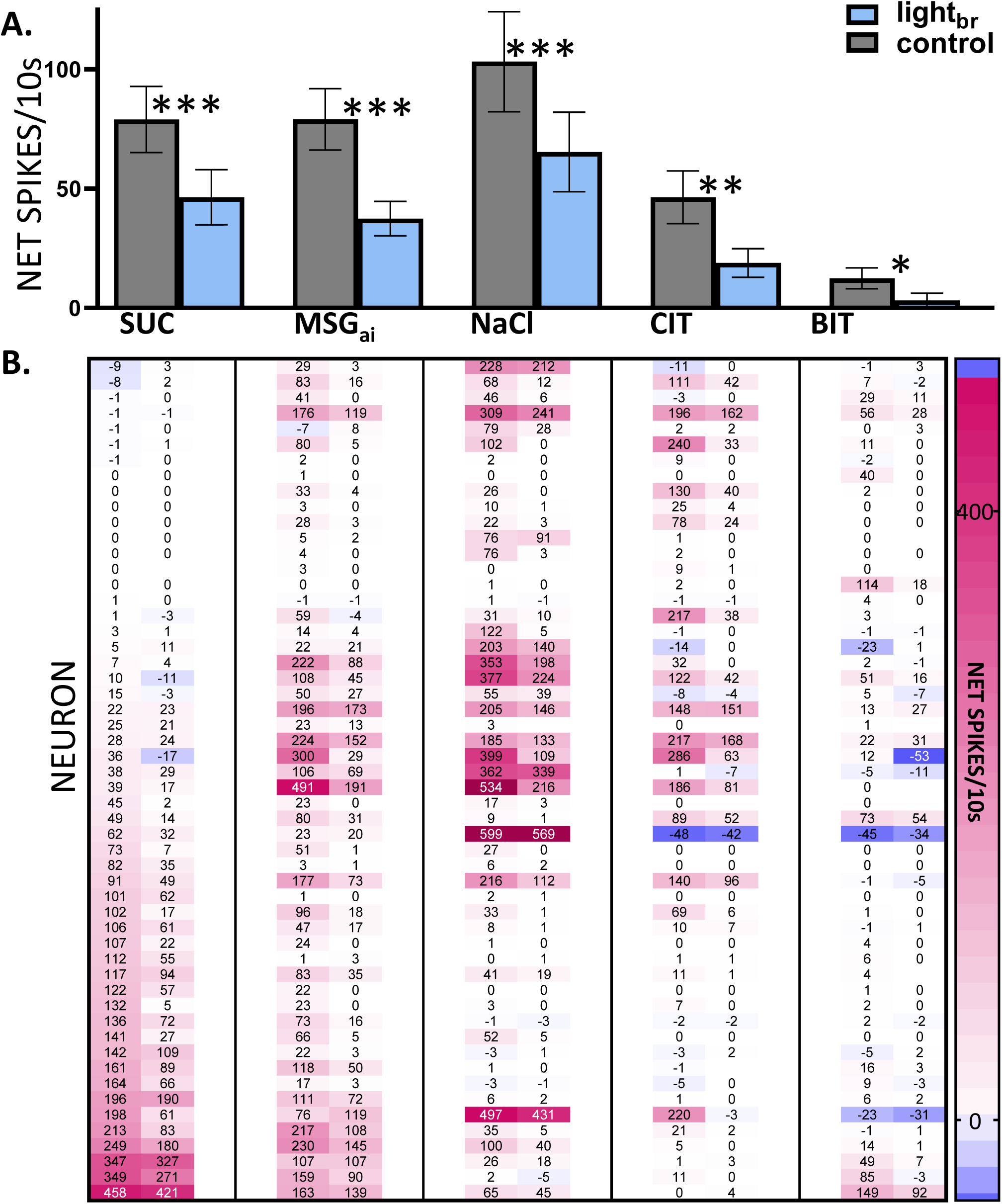
Responses to all taste qualities are suppressed by optogenetic stimulation of the GABA network. **A.** Mean net (± S.E.M.) responses to each stimulus for 54 G-_TASTE_ neurons tested with stimulus set A. A repeated-measures ANOVA showed main effects of light: (*p =* 5.11e-08) and stimulus: (*p =* 5.37e-05). (Because there were a few missing data points for the brain-light stimulated condition for stimuli ineffective under control conditions, the sample size for the ANOVA was reduced to 46). There was also a light X stimulus interaction (*p =*.004) but this was not indicative of differential effects of inhibition across stimuli: Brain light suppressed responses for each stimulus: Bonferroni-adjusted t-tests: SUC: \*\*\**p =*7.75e-07, N=54; MSG_ai_: \*\*\**p =*1.92e-05, N=53; NaCl: \*\*\*\**p =*1.53e-04, N=51, CIT: \*\**p =*.005, N=50, *BIT: *p =*.015, N=46). **B.** Corresponding heat map showing individual responses of each neuron and stimulus under light-stimulated and control conditions, lined up with the means in the panel A. Cells were ordered by their responsiveness to sucrose with no light. Response is color-coded according to the legend on the right and indicated by the numbers in a given rectangle; rectangles without numbers indicate missing data points.

#### Relative Response Profiles and Ensemble Coding Remain Stable During GABA Activation

Inhibitory effects for all qualities were likewise apparent when neurons were segregated according to chemosensitive profile. A cluster analysis of control responses suggested four main groups principally defined by the quality(ies) associated with the standard taste stimulus or stimuli that elicited the largest response(s) (**Figure 8A**). Although the cluster tree reveals heterogeneity within groups, the average response profiles for the groups were distinctive. Neurons responding optimally to sucrose also responded robustly to MSG_ai_, but poorly to other qualities and thus were labeled “SWEET/UMAMI” cells. Another chemosensitive cluster (“Na^+^”, N=14) responded 3X as well NaCl as any other stimulus including MSG_ai_, despite its higher sodium content, presumably reflecting attenuation by the amiloride in the MSG_ai_ cocktail. A third group responded nearly equivalently to NaCl, citric acid and MSG_ai_, and thus was considered an electrolyte generalist group (“EG”, N=13). The small number of remaining neurons were characterized by responding optimally and nearly exclusively to the bitter mixture (“BIT”, N=3). ANOVA indicated significant main effects of light and stimulus and an interaction between neuron group and stimulus but no interaction between light and neuron group, suggesting that all chemosensitive types were similarly susceptible to inhibition (see Figure caption for details). To probe this further, we analyzed the degree of absolute and proportional suppression for the classic taste stimulus eliciting the largest response in a given neuron; again, there was no impact of neuron group on this measure (t-tests, *p’s* >.1).

**Figure 8.**
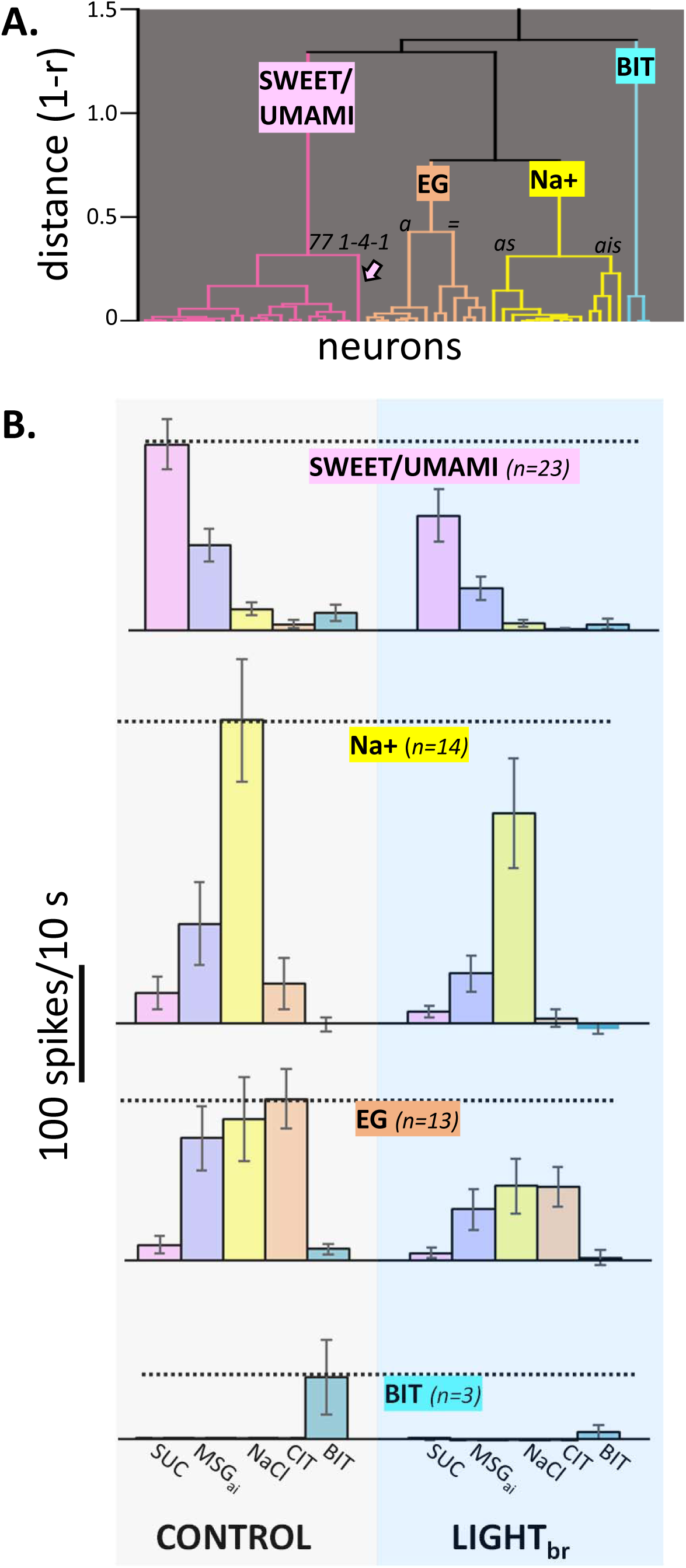
Activation of the GABA network suppresses responses in all chemosensitive clusters but the shape of the profiles is largely unchanged. **A**. Cluster tree showing segregation into 4 main chemosensitive groups. One neuron was omitted from the clustering process because it only joined the rest of the neurons at a distance (1-r) of 0.898. The resulting clusters largely reflected the stimulus or stimuli evoking the optimal response(s) in a cell: sucrose/MSG_ai_ (“SWEET/UMAMI”), NaCl (“Na+”), electrolyte generalists (“EG”, comparable mean responses to MSG_ai_, NaCl, and CIT), and a small group responding nearly exclusively to the quinine/cycloheximide cocktail (“BIT”). We used these 4 groups for further analysis to be parsimonious and to maintain adequate sample sizes. However, there was heterogeneity within the Na^+^ and EG groups: The “Na^+^” group was mostly comprised of cells with optimal responses to NaCl and minimal MSG_ai_ responses (“as”-amiloride sensitive, N=10). A smaller group of Na^+^ cells (“ais”-amiloride insensitive, N=4) responded optimally to NaCl but had robust responses to MSG_ai_. The EG group split into neurons that responded similarly to MSG_ai_, NaCl and CIT (N=6, “=”) and those with a more dominant response to CIT (N=7, “a”). The arrow indicates the last neuron that joined the “SWEET/UMAMI” cluster and is referred to in Figures 9 & 10. **B**. Mean (± S.E.M.) response profiles for each cluster under control conditions and during activation of the GABA network by light_br_. Activating the GABA network had a potent effect on response magnitude but the relative order of effectiveness of the stimuli is virtually identical under the two conditions. Mixed ANOVA, main effects-cluster: *p =*.229; light_br_: *p =*2.37e-05, and stimulus: *p =*1.95e-04, no interaction between cluster and light_br_ : *p* = .233. There was an interaction between stimulus and cluster (*p =*9.99e-16), suggesting that the neuron groups were distinct. The sample size for the ANOVA was reduced to 45 due to a few missing data points for the light condition for stimuli that did not evoke a response under control conditions. When the ANOVA was repeated with the full data set by substituting missing values with the control (null) values, conclusions were identical.

Importantly, though mean responses were clearly smaller during optogenetic inhibition, **Figure 8B** shows that the shapes of the average chemosensitive profiles were highly similar during GABA activation. The average order of effectiveness across stimuli was virtually identical under either condition. Ten cells (3 SWEET/UMAMI, 2 Na^+^, 3 EG and 2 BIT) became entirely unresponsive during optogenetic stimulation but the stimulus eliciting the largest response was unaltered for 40 of the 44 remaining neurons. Indeed, across individual cells, the correlation between response profiles for the two conditions was very high (mean r = +0.96 ± 0.13, N=44; **Figure 9A**), with all but one cell exhibiting a correlation greater than +0.85.

**Figure 9.**
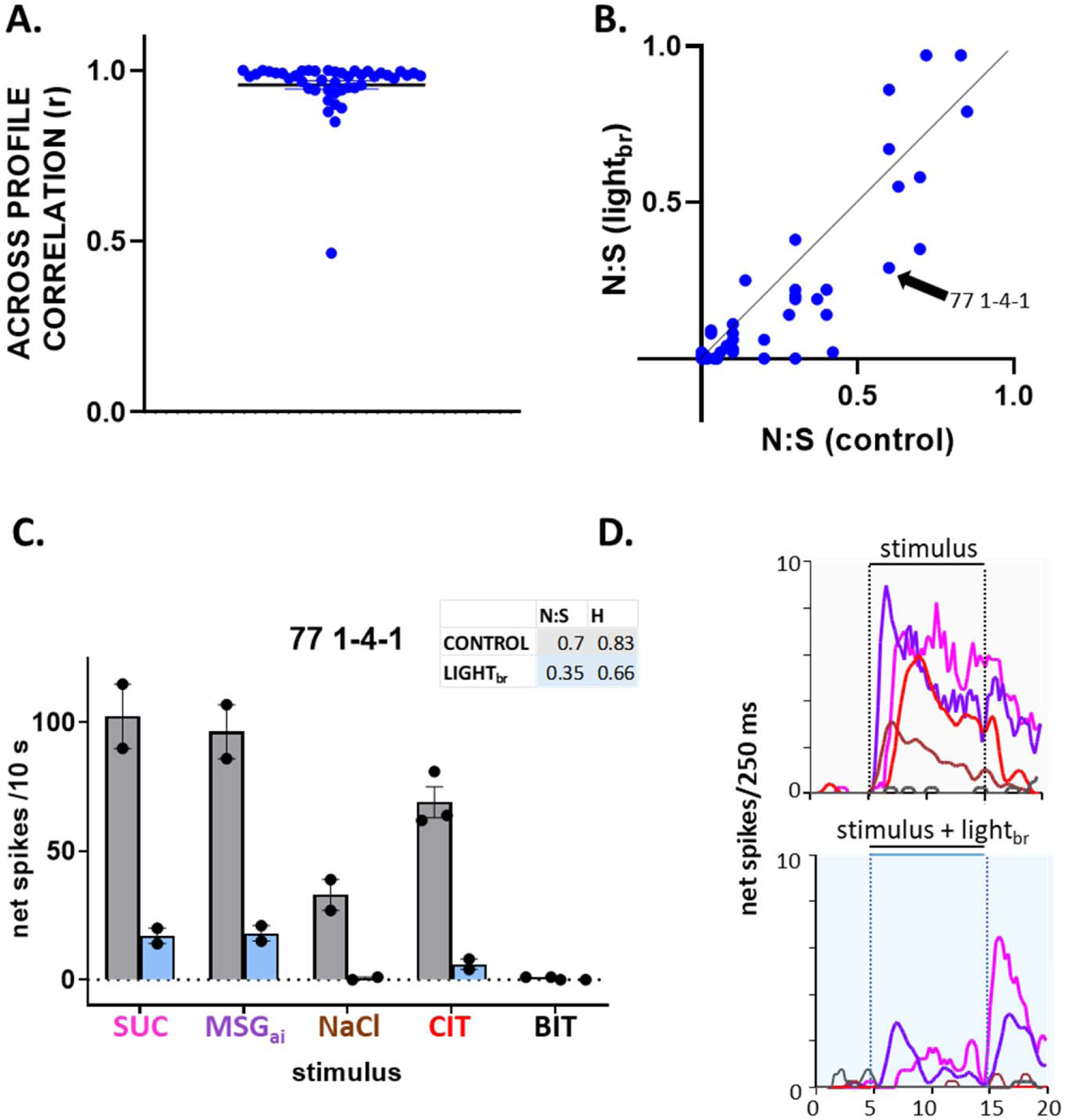
Response profiles are highly similar but modestly narrower under GABA activation. **A.** Across-profile correlations (Pearson’s r) for individual neurons during control conditions and during light_br_. Dots show results for individual cells; horizontal lines indicate mean and S.E.M. **B.** Scatterplot depicting the noise:signal ratio (N:S) during control conditions and during light_br_. Points on the solid line indicate a neuron with the same N:S ratio under the two conditions. Cells below the line have narrower tuning and those above have broader profiles with light. The arrow points to cell 77 1-4-1, illustrated in **C-D** that shows an example of sharpened tuning under GABA activation. **C.** Under control conditions, neuron 77 1-4-1 was very broadly tuned (Figure 8 shows that it was the last cell to join the SWEET/UMAMI cluster). This cell responded significantly to 4/5 qualities, reflected in high N:S and entropy measures. During light_br_, responses were greatly depressed and only SUC and MSG_ai_ remained effective. Moreover, both the N:S and entropy measures declined. Bars indicate mean 10 s net responses to the 5 stimuli; dots show individual replications; the vertical line indicates the S.E.M. **D.** Average time course (250 ms bins) of the responses. Responses are color-coded as in panel C. Interestingly, responses to MSG_ai_ and NaCl occurred at a shorter latency than SUC and CIT and the relative time course of SUC and MSG_ai_ remained unchanged under light_br_. Because responses to stimulation of both the anterior and posterior mouth were recorded on this track, we speculate that the latency differences may be due to differences in the receptive fields for these two stimuli.

Despite this high degree of correlation, neurons did become more narrowly tuned during optogenetic inhibition. These effects were statistically significant, albeit modest. For cells that remained responsive during light_br_ stimulation, we assessed tuning using three measures: (1) the number of responses meeting criteria, (2) the entropy measure and (3) the noise:signal (N:S) ratio. With each measure tuning became sharper during light_br_ stimulation. Under control conditions, the average number of responses in a cell was 2.8 ± 0.2 which decreased to 2.1 ± .2 during optogenetic stimulation (*p* = 4.54e-05, N=44). Likewise, entropy (H) declined from 0.537 ± 0.031 to 0.437 ± 0.039 (*p* = 4.53e-05, N=44) and the N:S ratio changed from 0.27 ± 0.04 to 0.20 ± 0.04 (**Figure 9B**, *p =*.002). An ANOVA incorporating chemosensitive group as a factor showed main effects of light_br_ for each measure (N:S, *p* = .003; number of responses: *p* = 5.14e-5; entropy, *p* = 2.61e-04) and a main effect of cluster for both entropy (*p* =.023) and N:S (*p =* 7.35e-05), but no interaction between chemosensitive group and light_br_ for any measure (N:S, *p* = .91; number of responses: *p =* .61; entropy, *p* =.33), suggesting a common effect in narrowing tuning across chemosensitive clusters. **Figure 9C&D** illustrates one of the more prominent effects of inhibition on tuning that occurred in a particularly broadly-tuned SWEET/UMAMI cell.

To assess whether these changes in tuning affected ensemble coding, we calculated across-neuron correlations and depicted them in a multidimensional scaling plot (**Figure 10**). Relationships among stimuli were strikingly similar for control responses and responses suppressed by stimulation of the GABA network. Although across-neuron correlations did exhibit some subtle shifts during optogenetic stimulation, these shifts were not sufficient to alter the overall pattern. Moreover, a given stimulus was positioned very closely in the taste space during the control and optogenetic stimulation conditions and correlations between these pairs were high (r = 0.72 - 0.93), exceeding correlations between any other pair of stimuli. Thus, we conclude that the main effect of non-selectively activating the rNST GABA network is to change response gain; in other words, the effects of inhibition are largely divisive rather than subtractive (Semyanov et al., 2004; Atallah et al., 2012; Wilson et al., 2012; Chen et al., 2016).

**Figure 10.**
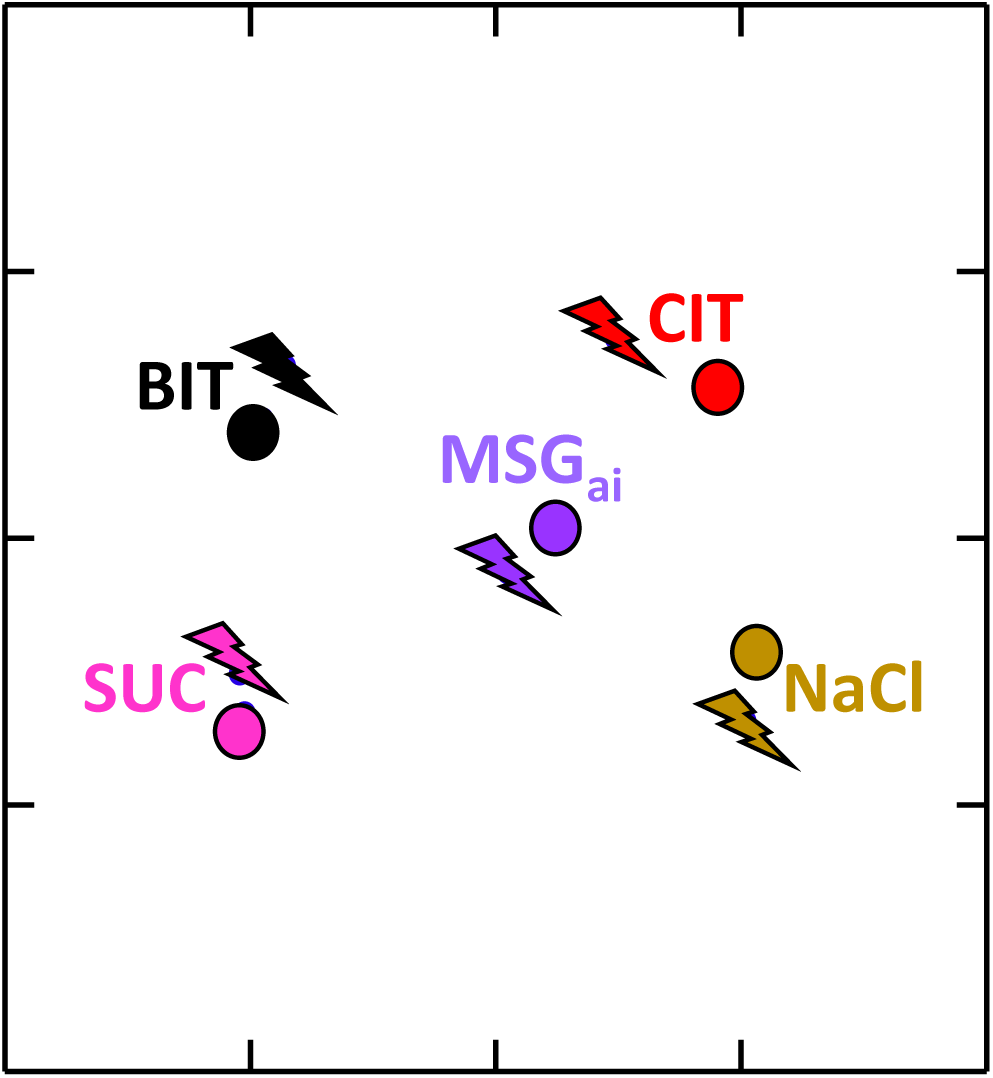
Ensemble patterns remain stable during GABA network activation. Multidimensional scaling plot of the 5 stimuli under control conditions (dots) and during optogenetic activation of the GABA network (lightning bolts). The placement of the stimuli is highly similar under the two conditions.

#### Effects of GABA Network activation on Mouth-Light Driven Responses

One limitation of the conclusion that activation of the inhibitory network mostly affects gain is that the present sample was dominated by SWEET/UMAMI and Na^+^ (amiloride-sensitive) neurons, types that remained narrowly tuned despite attempts to broaden profiles by using relatively high stimulus concentrations. To overcome this limitation, we used a second approach to activate cells over a more controlled range of firing rates by stimulating taste bud cells expressing ChR2 with blue light pulses directed to the mouth at different stimulation frequencies. We used this approach in a subset of G-_TASTE_ neurons when the cell remained well-isolated following taste stimulation. Indeed, a particularly high quality of isolation was necessary for the mouth-light protocol, since this stimulus often recruited additional small potentials that interfered with accurate quantification of light_m_-driven responses. Nevertheless, we were able to assess such responses in 16 neurons including 7 SWEET/UMAMI, 4 Na^+^ and 5 EG neurons. **Figure 11A&B** shows an example of the firing pattern and magnitude of light_m_ driven responses from a single neuron; Figure **11C** illustrates the mean responses from all 16 cells. Mean responses to light_m_ stimulation increased from ∼2-10 Hz but at higher stimulation frequencies responses saturated and then declined. Figure **11D** replots the mouth light responses in **11C** together with the taste responses from the same cells. In this graph, responses were ordered according to the efficacy of the largest control response for a given neuron, which yielded average “tuning curves” under both conditions. Tuning curves for taste responses exhibited a much sharper peak than did the responses to the range of mouth light frequencies tested, which declined more gradually.

**Figure 11.**
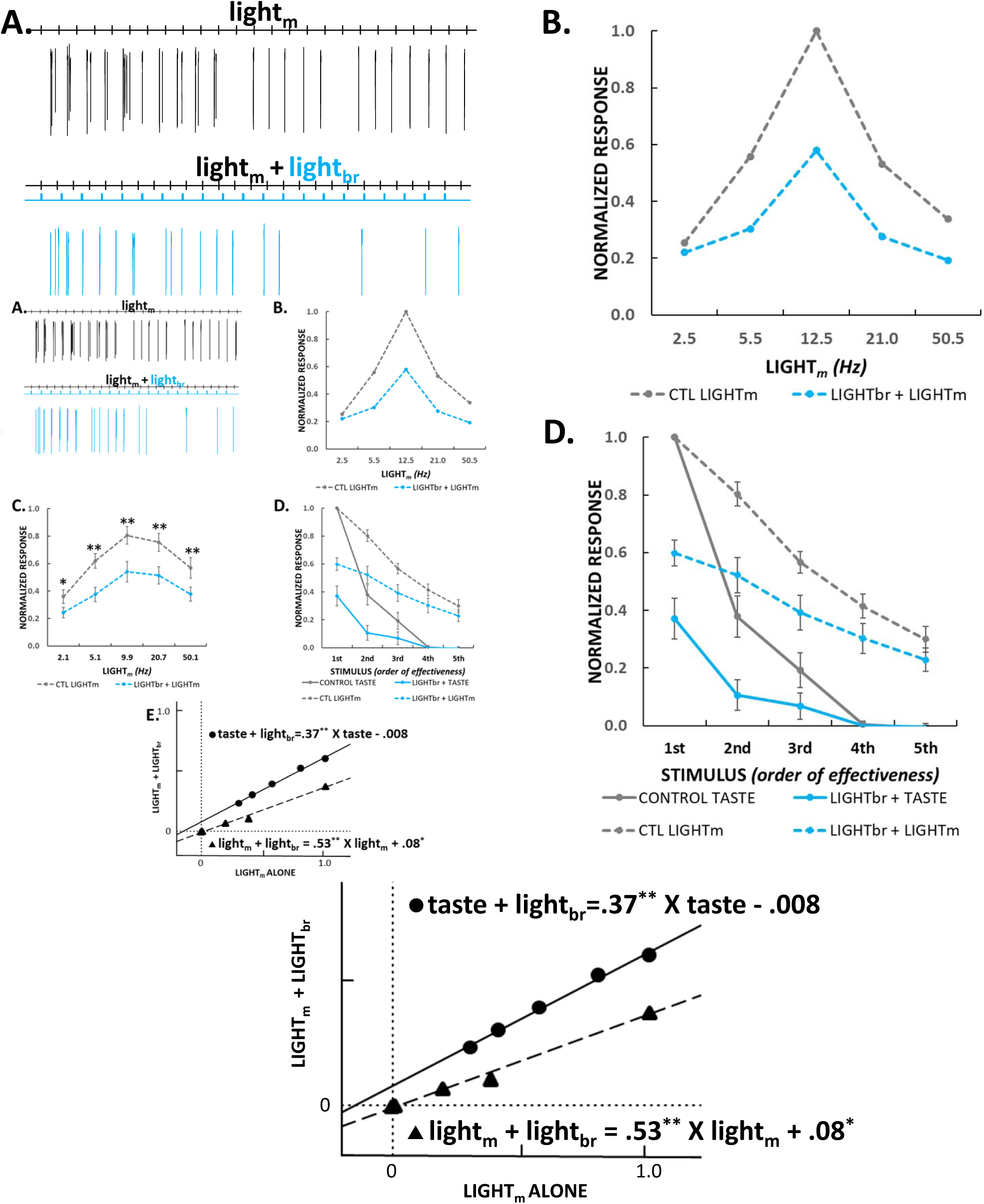
Threshold linear analysis of taste and light_m_ responses suggests a larger effect on gain than tuning. **A.** Firing pattern of an individual neuron to repetitive light_m_ stimulation (12.5Hz, 5ms, 10mW) under control conditions and during activation of the GABA network with light_br_ (10 Hz, 5ms). Light_m_ stimulation elicited time-locked firing that was suppressed by light_br_ stimulation. **B.** Responses from the cell shown in **A** at different stimulation frequencies. **C.** Average normalized response rates from stimulation at ∼2-50 Hz under control conditions and their suppression during light_br_. ANOVA: light, *p =* 2.84e-06, frequency, *p =* .041, light X frequency, *p =* .857. Bonferroni-adjusted t-tests: 2Hz, *p =*.015 (N=13); 5 Hz, *p =*.005 (N=16), 10 Hz, *p =* 1.18e-03 (N=16), 20 Hz, *p =* 2.11e-05 (N=16), 50 Hz, *p =*.005, (N=15). **D.** Tuning curves for 16 neurons tested with mouth light for light_m_ and taste responses derived by ordering normalized responses from largest to smallest under control conditions and then ordering stimuli during light_br_ identically. **E.** Linear regression of control versus light_br_ stimulation for taste and light_m_ responses. In both cases activating the GABA network had a larger effect on slope than on the Y intercept, suggesting a greater effect on gain than tuning.

We then applied linear regression to the relationship between the mean mouth-light or taste driven responses under control conditions and those suppressed by optogenetic stimulation of the NST GABA network (**Figure 11E**). In both cases, linear functions fit the average tuning curves well (r^2^ = 0.995 mouth- light; 0.991, taste) and the derived functions indicated a more prominent effect on slope (mouth-light: 0.533, *p =* 1.63e-04; taste: 0.371, *p =* 3.61e-04) than on the Y intercept (mouth-light: 0.08, *p =* .013; taste: -0.008, *p =*.458). Thus, for both mouth light and taste responses, the marked and significant deviation of the slope from unity, along with the small and, for taste, non-significant deviation in the intercept from zero, is consistent with the conclusion that inhibition is acting mainly in a divisive fashion.

### BEHAVIORAL EFFECTS OF ACTIVATING GABA NEURONS

Lastly, we activated GABA neurons in the rNST by expressing the excitatory DREADD, hM3D(Gq) in GAD65-expresing rNST neurons using viral injections and studied the behavioral effects on licking representative unpalatable (quinine) and palatable (sucrose and maltrin) stimuli. Injections were well-centered in the rNST (**Figure 12A**) and largely confined to the nucleus although some spread to the overlying vestibular and underlying reticular formation occurred. No CNO effect on sucrose licking was discernable in a control group with viral injections expressing mCherry alone (**Figure 12B**; see caption for all statistics). In contrast, licking to sucrose and maltrin decreased whereas licking to quinine was increased after CNO compared to saline injections (**Figure 12C-E**). This is consistent with what would be expected if neural responsiveness to these hedonically opposite stimuli were both suppressed. Fitting a logistic function to each of the mean curves demonstrated a rightward shift for CNO for each stimulus. It is notable that, in contrast to the neurophysiological effects that we observed, the behavioral effects are likely dependent mainly on local GABA neurons since this AAV virus is not transported retrogradely. Indeed, we did not observe labeling of neurons in the central nucleus of the amygdala (not shown).

**Figure 12.**
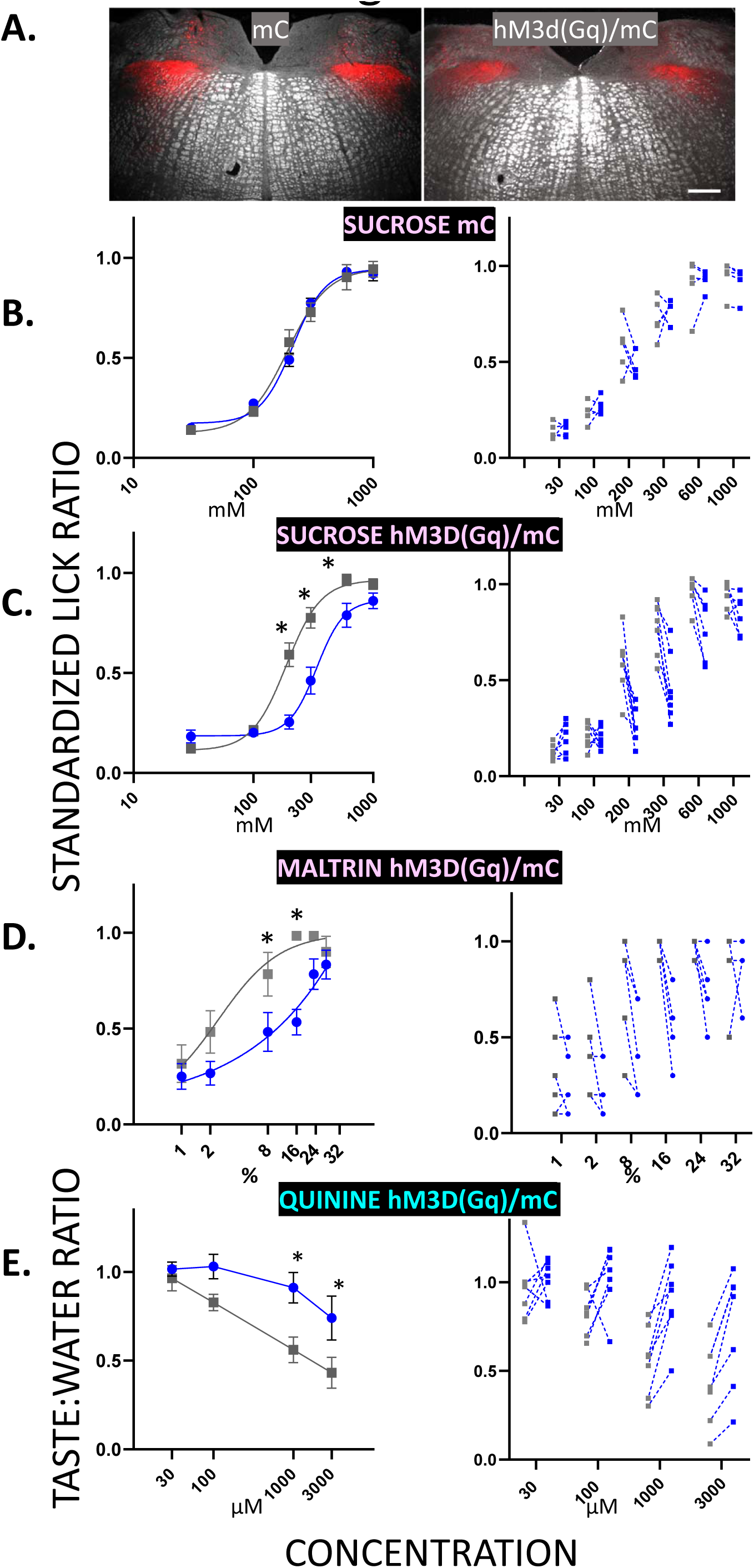
Activating rNST GAD65 neurons chemogenetically dampens behavioral acceptance of palatable (sucrose and maltrin) and rejection of unpalatable (quinine) and taste stimuli. **A.** Representative photomicrographs of mCherry (mC) expression in mice with injections of AAV2-hSyn-DIO-hM3D(Gq)-mcherry (mC) and AAV2-hSyn-DIO-hM3DGq/mcherry (hm3D(Gq)) in GAD65-cre mice. **B-F** Behavioral effects of CNO versus saline injections in mC (N=5) or hm3D(Gq)-injected mice (N=8). Symbols in the left panels show mean ± S.E.M. Asterisks indicate significance for Bonferroni-adjusted t-tests following ANOVA. Lines show curve fits for the equation: Y=Bottom + (Top-Bottom)/(1+10^((LogEC50-X)*HillSlope)^; an exception is quinine/CNO for the hm3D(Gq) mice, where the function could not be fit with this equation. Right panels show data points for individual mice. The same group of 8 mice was tested for all stimuli, but one or two mice were dropped from each analysis due to insufficient trials (dropped cases: GQ-13 for quinine, GQ-14 for sucrose and both GQ14 and 7 for maltrin); therefore, the final N’s for the three stimuli were 7 for quinine and sucrose and 6 for maltrin. **B**. CNO had no effect on sucrose licking in mice injected with a virus that only drove expression of mC (ANOVA, drug, *p =* .488, concentration, *p =* 9.99e-16, drug X concentration, *p =* .098; EC50 = 229mM, saline; 230mM, CNO. C. In contrast, CNO injection decreased standardized lick ratios for sucrose (ANOVA: drug, *p =*.007, concentration, *p =* 9.99e-16; drug X concentration: *p =* 3.47e-8. Bonferroni-adjusted t-tests: 200mM, \**p =* .021; 300 mM, *p =* .021 and 600 mM; *p =*.042; EC50 = 228mM saline; 252mM CNO and D. maltrin (ANOVA: drug, *p =*.007; concentration, *p =* 6.21e-10, drug X concentration, *p =*.004, Bonferroni-adjusted t-tests-significant* effects of CNO for 8%, *p =* .021 and 16%, *p =*.003. EC50 = 2.4% saline; 6.2 X 10^6^ % CNO). E. Activation of GAD65-expressing neurons increased quinine licking relative to water licks for the two highest concentrations (ANOVA: drug, *p =*.009, concentration, *p =* 7.23e-05, drug X concentration, *p =* .011. Bonferroni-adjusted t-tests: (1mM, \**p =* .028; 3mM, *p =*.012). We also observed a slowing of lick rate, as reflected in the modal session interlick interval (ILI) after CNO injection in the hm3D(Gq) group during the sucrose (138 ± 5 ms CNO vs 120 ± 2.7 ms, saline, *p =* .006) and maltrin (145 ± 5 CNO vs 119 ± 1 ms saline, *p =*.001) test sessions, but not during the quinine tests (123 ± 2 ms CNO vs 119 ± 1 ms saline, *p =* .093). There were no CNO effects on lick rate in the mC control group (112 ± 3 ms CNO vs 113 ± 3 ms saline, *p* > .1).

## Discussion

The present study is the first to describe gustatory response properties of identified rNST GABA taste neurons. Taste profiles of G+_TASTE_ neurons were similar to non-GABA taste neurons although response rates were dramatically weaker. Optotagging also afforded identification of GABA neurons unresponsive to taste stimulation. These three distinctive cell types were distributed non-homogeneously in the nucleus from dorsal to ventral. Activating NST extrinsic and intrinsic inhibitory circuitry markedly suppressed taste responses of G-_TASTE_ neurons, impacting all qualities and types of chemosensitive neurons. Nearly 20% were totally silenced. Behavioral effects of activating GABA neurons likewise included both palatable and unpalatable taste qualities. Using both traditional taste and optogenetic stimulation of taste bud cells to induce an afferent response allowed us to quantify the impact of centrally released GABA on response magnitude and tuning of G-_TASTE_ neurons. Using both approaches, GABA activation had a strong suppressive effect and modestly sharpened tuning curves of G-_TASTE_ neurons. These effects did not substantially modify the ensemble pattern for taste quality.

### Properties of rNST GABA neurons

A large group of rNST GABA neurons has been documented using immunohistochemistry (Lasiter and Kachele, 1988; Davis, 1993; Wang and Bradley, 2010), *in situ* hybridization (Stornetta et al., 2002) and multiple reporter lines (Wang and Bradley, 2010; Boxwell et al., 2013; Travers et al., 2018). There are likely subclasses of rNST inhibitory neurons as in other CNS regions. In any case, the GAD65 cre X ChR2 mouse we used likely encompassed a majority of GABAergic rNST cells based on the high degree of co-localization between GAD65 and VGAT in the nucleus (Travers et al., 2018) and comparable distributions of GAD65 and GAD67 expression in this region (Stornetta et al., 2002).

The G+_TASTE_ GABA taste neurons we encountered had less robust taste responses than G-_TASTE_ cells, consistent with their more ventral location, which would be expected to put them in sparser contact with the more dorsal gustatory primary afferents. This observation also agrees with previous *in vitro* experiments demonstrating fewer spikes in response to solitary tract stimulation, a smaller paired-pulse ratio, and saturation of firing rate at a lower level of depolarizing current in GABA than non-GABA neurons (Chen et al., 2016; Boxwell et al., 2018). The weaker responses of GABA taste neurons is also consistent with antidromic stimulation experiments demonstrating that NST taste neurons that do not project to the PBN are less responsive (Ogawa et al., 1984; Monroe and Di Lorenzo, 1995; Cho et al., 2002; Geran and Travers, 2009), and suggests that some non-PBN projection neurons in these studies were GABAergic. The modest number of G+_TASTE_ neurons we characterized prevented firm conclusions about more variable differences reported in antidromic studies including a narrower breadth of tuning (Monroe and Di Lorenzo, 1995; Geran and Travers, 2009) and a propensity for more non-projection neurons to respond to aversive stimuli (Cho et al., 2002; Geran and Travers, 2009). Each taste quality activated some G+_TASTE_ (and G-_TASTE_) cells, suggesting that inhibitory neurons are not preferentially devoted to processing specific tastes. On the other hand, it is perhaps notable that the one exception to lower evoked response rates for G+_TASTE_ cells was the bitter mixture, although this observation is tentative due to small sample of bitter-responsive G-_TASTE_ neurons in this study. In addition to gustatory-responsive GABA cells, a comparable population of rNST GABA neurons were unresponsive to taste or somatosensory stimulation (G+_UNR_). On average, these were ventral to G+_TASTE_ cells. The ventral bias for GABA cells (in general) is interesting since this region preferentially projects to the reticular formation (Travers, 1988; Beckman and Whitehead, 1991; Halsell et al., 1996; Zaidi et al., 2008). This suggests that the rNST inhibitory network has strong influence on modulating autonomic and oromotor reflexes, consistent with our observations that activation of NST GABA neurons by DREADDS can influence lick rate. It may also be functionally significant that some extrinsic pathways targeting rNST, e.g., the caudal NST (Travers et al., 2018) and central nucleus of the amygdala, preferentially target the ventral subdivision [reviewed in (Travers and Spector, 2021)].

rNST neurons unresponsive to orosensory stimulation have seldom been reported, at least in anesthetized preparations. This is not surprising since the G+_UNR_ we encountered had negligible spontaneous activity and were often identified only when optogenetically activated. It seems likely that these G+_UNR_ cells receive other central inputs. Indeed, Smith and Li (Li and Smith, 1997) showed that suppression of rNST taste responses by electrical stimulation of gustatory cortex could be abrogated by locally infusing a GABA_A_ receptor antagonist, suggesting an excitatory pathway from taste cortex to NST GABA neurons that in turn influence taste-responsive cells. In agreement, a recent study used monosynaptic rabies tracing and optogenetic activation of the insular-NST pathway to demonstrate largely excitatory connections to rNST somatostatin neurons (Jin et al., 2021) some of which are likely GABAergic (Wang and Bradley, 2010; Thek et al., 2019; Kalyanasundar et al., 2022). The caudal NST (Travers et al., 2018) and central nucleus of the amygdala also project to rNST although whether they make direct connections to GABA neurons is unknown (Saha et al., 2000; Saha et al., 2002; Bartonjo and Lundy, 2020; Jin et al., 2021). Interestingly, some studies in the awake rat report that many rNST cells are not taste responsive, but rather lick-rhythmic (Denman et al., 2019) but other studies did not encounter this population (Nakamura and Norgren, 1991, 1993).

### Effects of Activating the rNST GABA Network

Optgenetically activating rNST GABA neurons and fibers markedly reduced gustatory responses of G-_TASTE_ cells. Because ChR2 was globally expressed, our optogenetic stimulation targeted not only local GABA cells, but also inhibitory projections from other structures, including the central nucleus of the amygdala (Saha et al., 2000; Saha et al., 2002; Bartonjo and Lundy, 2020; Jin et al., 2021) and caudal NST (Travers et al., 2018). The affected non-GABA taste cells likely included glutamatergic neurons that project to PBN (Gill et al., 1999), which provides parallel inputs to the thalamocortical pathway as well as limbic structures (Norgren and Leonard, 1973) [reviewed in (Travers and Spector, 2021)]. Thus, NST neurons that ultimately contribute to both perceptual and motivational functions appear susceptible to inhibition. In addition, the rNST projects to the underlying reticular formation and caudal NST (Travers, 1988; Beckman and Whitehead, 1991; Halsell et al., 1996; Zaidi et al., 2008; Travers et al., 2018), substrates for coordinating ingestion and rejection oromotor responses (Chen et al., 2001) and visceral processing. Thus, inhibitory modulation likewise has potential to directly affect taste-elicited consummatory behaviors and signals from internal organs. Though our results demonstrate potential for GABAergic modulation of all taste qualities, more specific effects are likely in real-world situations. Two recent studies used optogenetics to suggest that amygdalar GABA projections comprise a substrate for bitter-induced suppression of sweet signals in rNST (Jin et al., 2021) or tonically inhibit bitter signaling (Bartonjo et al., 2022).

In agreement with *in vitro* studies in hamster and rat demonstrating that GABA or GABA agonists (Liu et al., 1993; Wang and Bradley, 1993) cause decreases in firing or increases in membrane resistance and hyperpolarization in most rNST neurons, we observed overwhelmingly suppressive effects upon activating the GABA network. Also consistent with the current findings, an *in vivo* study in the hamster did not report GABA-induced increases in spontaneous or taste-driven activity (Smith and Li, 1998). On the other hand, the hamster *in vitro* study did demonstrate that GABA infusion increased spontaneous rate in 11% of neurons (Liu et al., 1993) and a recent report observed heterogeneous increases and decreases in NST taste responses using optogenetic activation of virally-driven ChR2 in GAD67 neurons (Sammons et al., 2021). Thus, it seems likely that strong GABA activation in the current study masked less prevalent GABAergic disinhibitory circuits.

In addition to suppression, there was a modest sharpening of response profiles, consistent with broadening of rNST taste profiles when GABA_A_ antagonists were infused in vivo (Smith and Li, 1998). However, it was more impressive than the shapes of chemosensitive profiles were highly similar under inhibitory influences. Thus, effects of activating the rNST GABA network were largely divisive, not subtractive. This conclusion was bolstered by manipulating firing rate using optogenetic activation of taste bud cells expressing ChR2 to overcome the limitation that many cells in the current study were narrowly tuned to natural taste stimulation. These limited effects on tuning suggest that taste quality representation was largely unchanged, but that intensity was dampened.

The stable neurophysiological representation of taste quality under GABAergic challenge is further supported by the behavioral effects of activating the GABA rNST neurons, where appropriate but dampened acceptance or rejection of sucrose, maltrin, and quinine occurred (**Figure 12**). In parallel experiments using an inhibitory DREADD, sucrose and quinine licking curves shifted to the left when GABA neurons were inhibited but the stimuli still elicited appropriate behaviors: sucrose preference and quinine rejection (Travers et al., 2020). We speculate that the GABA network permits a faithful, although amplified or dampened quality message to be transmitted by the first-order gustatory relay to permit adaptive adjustments in behavioral responses to taste stimuli under different homeostatic states and as a function of experience.

## Acknowledgements

This work was supported by NIH DC00416 to S.P.T. The excellent technical assistance of Cemaliye Semmedi, Sophia Pilloli, Jacob Harley, and Andrew Harley are greatly appreciated.

---

**Extended Data Figure 1-1.**
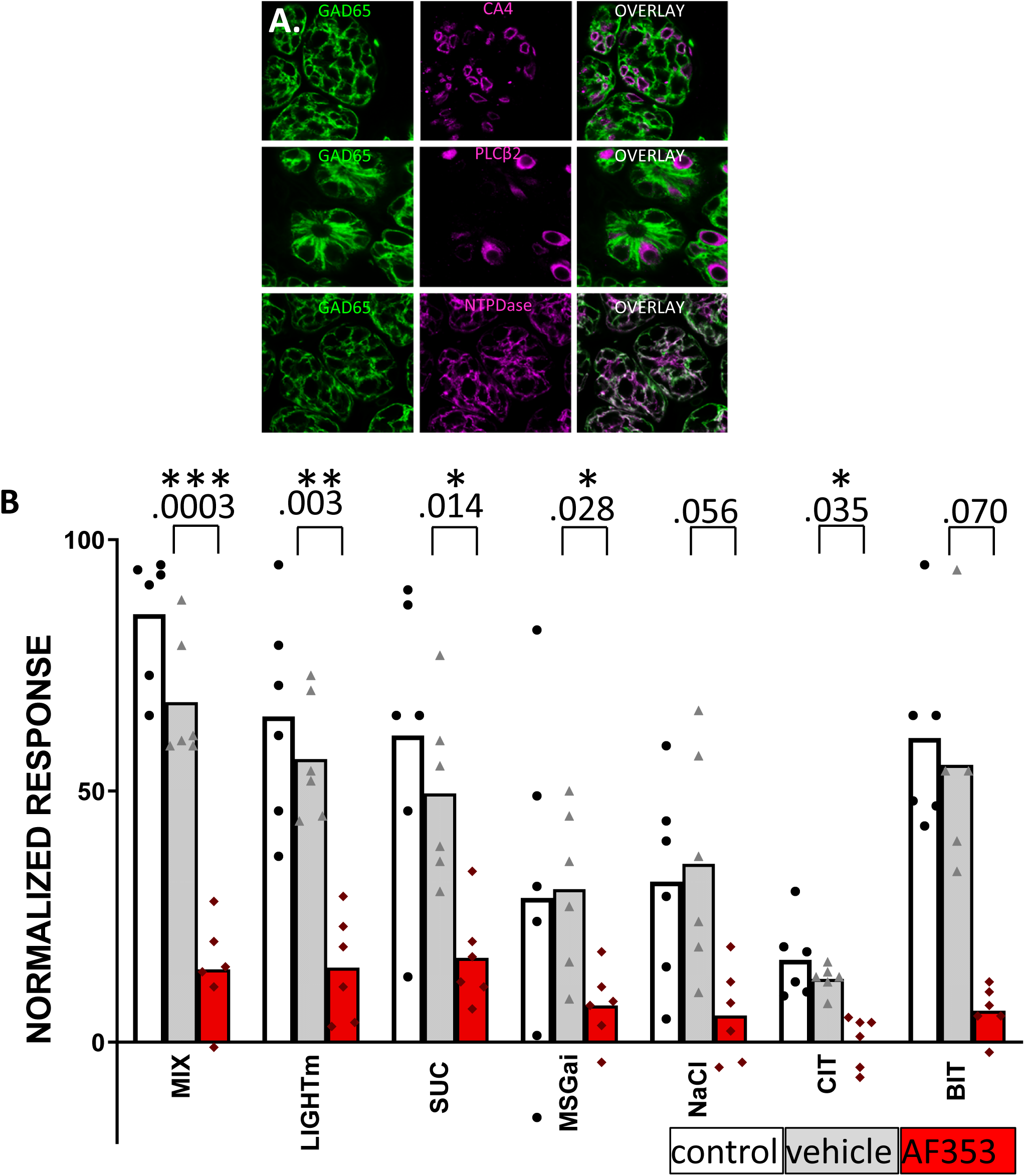
Most taste bud cells in the GAD65cre X ChR2/EYFP cross are Type I cells. Activating these taste bud cells optogenetically evokes responses in NST taste neurons that are dampened by lingual application of a P2X3 antagonist. **A. Immunostaining of taste buds from the GAD65cre X ChR2/EYFP cross.** Mice were perfused with phosphate-buffered saline followed by paraformaldehyde/lysine-metaperiodate and 40 µm frozen sections of fungiform and circumvallate papillae cut on a sliding microtome. Standard double-labeling immunofluorescent techniques with appropriate antibodies were used to identify Type I (NTPDase -Sevigny, host: rabbit, RRID: AB_2314986, 1:1000), Type II (PLCβ2 - Santa Cruz, host: rabbit, RRID: AB_2314986, 1:50 and Type III cells (carbonic anhydrase 4 [CA4] - R&D Systems, host: goat, RRID: AB_2070332, 1:1000). Analysis was performed inspecting 1 µm confocal z-stacks for each of the three stains (PLCβ2 - 52 cells from 2 fungiform & 8 circumvallate papillae; CA4 - 60 cells from 4 fungiform & 6 circumvallate papillae; NTPDase - 3 fungiform & 7 circumvallate buds - individual cells not counted. None of the PLCβ2- or CA4-labeled cells co-expressed EYFP. In contrast, there was extensive co-labeling of taste bud cells stained for NTPDase suggesting that most ChR2/EYFP-expressing cells in this mouse line are Type I cells (Baumer-Harrison et al., 2020; Larson et al., 2021; Rodriguez et al., 2021). Scale bar= 10 µm. **B. Ionotropic P2X3 receptors are involved in conveying mouth-lite driven responses centrally to a similar degree as taste responses.** Multiunit NST recordings from the same strain of mouse as in the main part of the study. The data is from a separate series of experiments using similar techniques to record multiunit responses to probe the role of ionotropic P2X3 receptors in conveying light_m_ responses centrally. Ionotropic P2X2/P2X3 heterodimers (and some homodimers) are present on primary afferent taste neurons and responsible for transmitting responses from taste buds to the primary afferent nerves (Finger et al., 2005). The marked suppression of responses to both light_m_ and taste stimuli after topical application of a P2X3 antagonist suggests a common mechanism in conveying responses to the primary afferents, although this effect could also reflect crucial signaling within the bud (Rodriguez et al., 2021). Bar graphs depict the mean magnitude of multiunit responses to a taste mixture, individual tastants and light directed at the oral cavity (5 ms/10hz/10mW) through a 400 µm optical fiber prior to (control) and after delivery of a vehicle (polyethylene glycol- PG, 10%) and a P2X3 antagonist (AF353, 1 mM dissolved 10% PG, (Vandenbeuch et al., 2015). Data are from 6 mice (individual mice denoted by dots) and responses were normalized to the maximum control response. Durations of PG and PG + AF353 application were 18±1 and 20±1 minutes. Stimulus abbreviations and concentrations: SUC: sucrose (300mM), MSG_ai_: (100mM MSG, 2.5mM IMP and 100um amiloride), NaCl: (100mM), CIT: citric acid (10mM), and BIT: a bitter cocktail of cycloheximide, (10uM) and quinine monohydrochloride (2.7mM). ANOVA: significant effects for condition (p = 5.55e-06), stimulus (p = 2.73e-06) and a condition X stimulus interaction (p = 8.77e-05). Pairwise ANOVA comparisons between conditions indicated significant differences between the control and post-AF (p =.001) and vehicle vs post-AF (p = 8.5e-05) but not between the control and vehicle conditions (p =.803). ANOVAs were done without responses to the bitter cocktail due to a missing data point for the post-AF condition. Significant differences between post-PG (vehicle) and post-AF conditions based on Bonferroni-adjusted t-tests are indicated by asterisks and p values above the bars. Responses to the bitter stimulus were analyzed separately based on 5 instead of 6 cases.

